# Attention Across Scales: From Individual Variation to Social Hierarchies and Brain Networks in Semi-Free-Ranging Macaques

**DOI:** 10.64898/2026.09.21.752444

**Authors:** Simona Vaitekunaite, Sarah Silvère, Hélène Meunier, Sébastien Ballesta, Ilaria Sani

## Abstract

Attention is a fundamental brain function supporting perception, decision-making, and social behavior, and its dysfunction profoundly impairs daily life. It is both dynamic and stable, varying across observations and individuals, changing across the lifespan, and being shaped by social and environmental experience. Yet capturing this complexity remains a central challenge in neuroscience. Here, we integrated longitudinal behavioral assessments of semi-free-ranging macaques living in naturalistic social groups with resting-state fMRI. We quantified performance across days, ages, and social hierarchies and related it to intrinsic brain organization. Distinct attentional phenotypes emerged, including individuals with reduced attentional control. Performance followed an inverted-U lifespan trajectory, improving from childhood to adulthood before declining. Social status modulated attentional performance. Critically, nonlinear lifespan trajectories and associations with individual attentional differences were most clearly expressed in frontoparietal connectivity. Together, these findings reveal how sustained attention is organized across scales, providing a biological framework for its individual diversity, social modulation, and neural basis.

## Introduction

Attention is a core organizing principle of adaptive behavior. Across human and nonhuman primate societies, it shapes how individuals detect, prioritize, and act on information in complex, dynamic environments. Among its forms, sustained attention—the capacity to maintain goal-directed engagement over time—is particularly critical for monitoring infrequent events, resisting distraction, and navigating socially structured worlds^1,2^.

Yet attention is inherently dynamic. It fluctuates across moments^3^, reflects stable inter-individual phenotypes^4,5^, evolves across the lifespan^4^, and varies with social and environmental contexts^2,6^. Disentangling these dynamic and trait-like components—and identifying their neural substrates—remains a central challenge in neuroscience. Human studies are often cross-sectional^7^, constrained by longevity, and confounded by educational, cultural, and socioeconomic variability^8^. As a result, it remains difficult to determine how attentional dynamics unfold over time and across contexts, and how they relate to intrinsic brain network organization.

Non-human primates offer a unique opportunity to overcome these limitations. Their biological proximity to humans^9–11^, shorter lifespans^12,13^, and richly structured social organizations^14,15^ allow cognitive trajectories to be tracked repeatedly, within and across individuals, in their natural environment, and over meaningful portions of the lifespan. However, traditional laboratory settings—characterized by very small group sizes and restricted environments—limit ecological validity and attenuate the natural social demands that shape attentional function^16,17^. Despite decades of research, attention has rarely been examined as a multi-level cognitive function that evolves within structured groups. Most work isolates individuals from their ecological and social contexts, implicitly treating attention as a laboratory-bound cognitive capacity rather than a dynamic function embedded in real-world systems. As a result, we lack an integrated account of how attentional abilities emerge, stabilize, and diverge across individuals living in natural hierarchies—especially given that attention operates across multiple *scales*.

These multiple scales encompass distinct sources of variation. At the shortest timescale, performance can fluctuate across moments, sessions, and days (*state fluctuations*), reflecting transient changes in fatigue, stress, or motivation^18^. Across longer periods, individuals can nevertheless maintain reproducible profiles of attentional performance (*stable phenotypes*)^1^, with some consistently exhibiting higher or lower levels of sustained attention across contexts, both within the range of normative variability^4^ and in clinical conditions such as attention-deficit/hyperactivity disorder (ADHD)^19^. Attentional capacities also change systematically with age (*lifespan trajectories*) and are shaped by social and environmental context (*social scale*). In humans, sustained attention follows a nonlinear trajectory^4^—improving through childhood and adolescence, peaking in adulthood, and declining in older age. Finally, these behavioral dimensions may be reflected in the organization of distributed brain networks (*neural scale*).

Among these distributed brain systems, the frontoparietal network, encompassing interconnected parietal and prefrontal cortices, such as the lateral intraparietal area (LIP) and the frontal eye field (FEF), plays a central role in sustaining goal-directed behavior over time^20–22^. Disruption, disconnection, or atypical development of this network has been implicated in ADHD^23^ and in age-related attentional decline^24^, suggesting that both phenotyping and lifespan changes in attention may be reflected in frontoparietal circuit organization. Crucially, its structural and functional conservation across primates^25–29^ suggests that core mechanisms of attentional control are deeply preserved. If sustained attention reflects a conserved control system, then developmental improvement, adult stability, and later decline should be mirrored in corresponding trajectories of frontoparietal connectivity. Moreover, stable inter-individual differences in attention should be embedded within this network’s intrinsic functional architecture rather than in purely visual systems. Although the frontoparietal network is structurally and functionally conserved across primates, how its connectivity is organized across these dimensions remains poorly understood.

Here, we integrated longitudinal behavioral and cognitive assessments within a population of semi-free-ranging Tonkean macaques (*Macaca tonkeana*) living in a cohesive social group with access to automated touchscreen testing^30,31^. This ecologically grounded setting enabled repeated assessment of sustained attention across developmental stages while preserving naturalistic social dynamics. We measured attentional performance using a well-established cross-species analogue of the human Continuous Performance Task^32^, quantified hierarchical status^31^, and characterized intrinsic brain organization using resting-state fMRI.

We show that distinct and stable attentional phenotypes emerge within the population and that sustained attention follows a nonlinear lifespan trajectory in macaques, mirroring human development and being further modulated by social hierarchy. Frontoparietal functional connectivity most clearly tracks both developmental change and individual differences.

Together, these findings reveal how sustained attention is organized across scales, linking behavior to conserved brain networks and providing a biological framework for understanding its evolution and individual diversity.

## Results

### Distinct and Stable Attentional Phenotypes Emerge Within the Population

We quantified sustained attention in a population of 32 semi-free-ranging *Macaca tonkeana* living in a socially structured, naturalistic environment (**Fig. 1a**). Animals had unrestricted access to automated touchscreen devices for cognitive testing and enrichment (**Fig. 1b**), enabling voluntary, repeated task engagement over extended periods. Following task acquisition, individuals were monitored for an average of 49 months, completing up to 7’973 sessions (**Fig. S1**). Social interactions were continuously tracked to derive hierarchical metrics (see Methods). A group of animals underwent resting-state functional MRI (rs-fMRI; n=18) scanning, and a smaller subset contributed both behavioral and imaging data, enabling direct brain–behavior comparisons (n=8). This innovative research strategy yielded dense longitudinal behavioral sampling across developmental stages alongside intrinsic functional connectivity measures.

**Figure 1.**
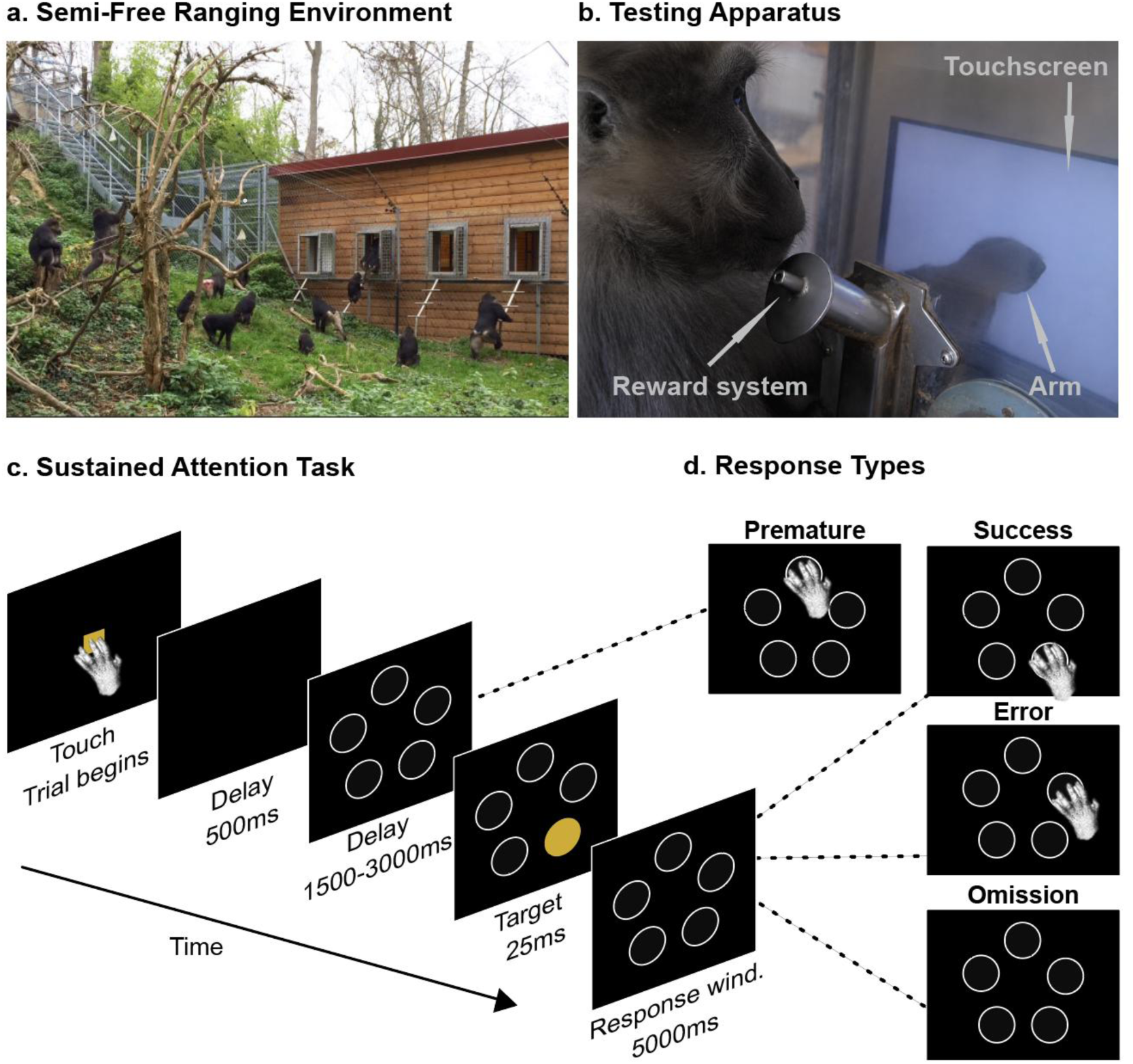
Experimental Subjects, Testing Apparatus, and Task Overview. **(a)** Semi-free-ranging macaques (M. tonkeana) in naturalistic outdoor enclosures with up to four touchscreen devices integrated into a wooden structure. **(b)** Interior view of a monkey interacting with a touchscreen connected to a fluid reward delivery system. **(c)** Schematic overview of the 5-Choice Serial Reaction Time Task, assessing sustained attention and inhibitory control through detection of a briefly flashed target. **(d)** Each trial yielded four possible outcomes: success (i.e., correct selection of the target location); premature response (i.e., selection before target onset); error (i.e., selection of a non-target location); and omission (i.e., failure to respond within the response window).

To assess sustained attention under ecologically grounded conditions, we employed a touchscreen-based adaptation of the 5-Choice Serial Reaction Time Task^32,33^ (**Fig. 1c**; see also Methods). Animals monitored multiple spatial locations and responded to a brief, unpredictable target flash while withholding premature responses. Each trial yielded four possible outcomes (**Fig. 1d**): successful target detection, premature responses, incorrect target selection, or omissions. Average task performance was high, and variability in success rates was driven predominantly by premature responses rather than errors in target selection (**Fig. 2a**), indicating that attentional differences primarily reflected inhibitory control demands rather than perceptual failures.

**Figure 2.**
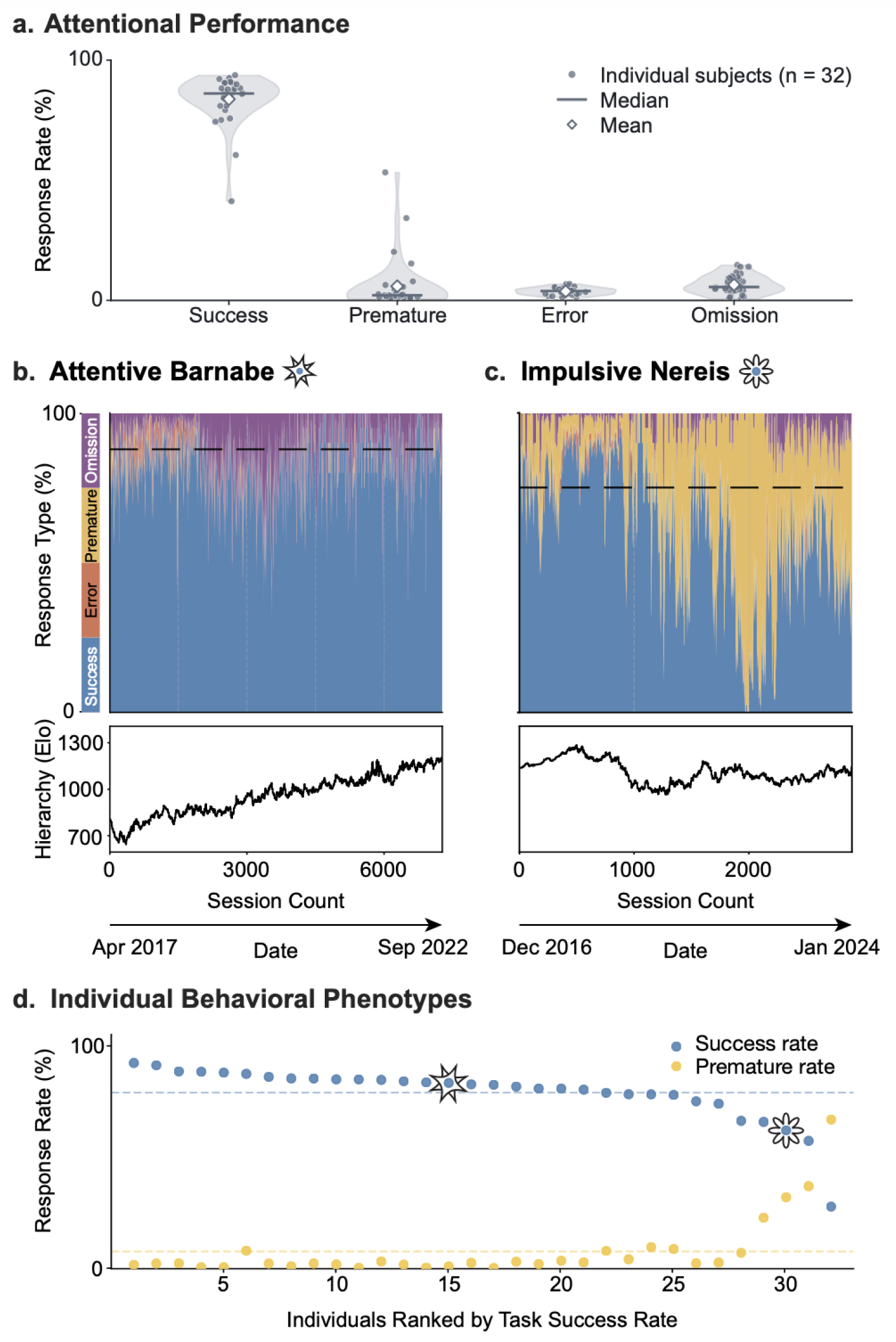
Overall Performance and Longitudinal Phenotypes. **(a)** Overall attentional performance across individuals. Violin plots show the distribution of individual response rates for each response type. **(b-c)** Longitudinal session-level trajectories for two individuals illustrating distinct behavioral phenotypes. Stacked area plots show the rolling proportions across 10-session windows for successes (blue), errors (orange), premature responses (yellow), and omissions (purple). The horizontal dashed line indicates each individual’s overall mean success rate, averaged across sessions. Lower panels show corresponding Elo rating trajectories over the same period, reflecting changes in social dominance rank. Elo ratings were initialized at 1000; increasing values indicate rising social rank, whereas decreasing values indicate declining social rank (see methods). **(d)** Mean success (blue) and premature (yellow) response rate for all 32 individuals, averaged across sessions, ranked from highest to lowest mean success rate. Dashed lines indicate the group means for each measure; star markers identify the representative individuals shown in b-c.

To determine whether attentional differences reflect stable biological traits rather than transient fluctuations, we examined longitudinal session-by-session behavioral trajectories over extended periods of time (**Fig. S1**). This analysis revealed distinct behavioral phenotypes that showed substantial stability over several years of observation. Most individuals displayed consistently high proportions of successful responses, with relatively few errors, omissions, or premature responses (**Fig. 2b**; **Fig. S2**). Among the remaining individuals, some exhibited transient periods of elevated premature responding against a background of otherwise good performance, whereas others showed chronically elevated premature responding across most sessions, accompanied by comparatively fewer successful responses (**Fig. 2c**; **Fig. S2**). We confirmed the temporal stability of these phenotypes quantitatively across successive testing windows, with high temporal consistency for both success rates and premature responding (intraclass correlation: success = 0.70; premature responding = 0.79; see Methods).

The phenotype showed no robust relationship with visual errors but generalized to premature responding in two independent tasks. In a visual control task where animals identified a small visual target on the touchscreen (**Fig. S3a**; see also Methods), performance was markedly different with virtually absent premature responding due to a shorter fixed delay, and interindividual variation manifesting primarily as errors (**Fig. S3c,e**). Visual-task error rates in this task were not associated with the sustained attention task error rates (Spearman’s ρ = 0.42, n = 18, p = .086, Bonferroni-adjusted α = .025), arguing against basic visual impairment as the primary explanation for the attentional phenotype. Interestingly, premature responding generalized to an independent cognitive control task, a delayed matching-to-sample paradigm that required short-term maintenance of object identity before selecting the target among distractors (**Fig. S3b**; see also Methods): individuals with elevated premature responding in the sustained attention task also tended to exhibit higher premature responding in this task (**Fig. S3d,f**; Spearman’s ρ = 0.64, n = 32, uncorrected p < .001, Bonferroni-adjusted α = .025), consistent with a broader trait-like dimension of attentional control. Notably, changes in attentional performance appeared to track social status in some individuals (**Fig. 2c**, bottom panel), suggesting a potential link between attention and social hierarchy that we examined formally in subsequent analyses.

Taken together, these findings reveal that sustained attention is not homogeneously distributed across individuals but is organized into stable phenotypic dimensions that persist over extended periods of observation.

### Sustained Attention Follows an Inverted-U Trajectory Across the Lifespan

While stable attentional phenotypes capture enduring differences between individuals, attentional abilities also change across development and aging. We therefore asked whether sustained attention follows a characteristic lifespan trajectory in this primate population. Behavioral performance was initially modeled as a function of age using linear and quadratic regressions of each individual’s average performance, allowing us to distinguish continuous monotonic age-related changes from the predicted inverted-U developmental profile (**Fig. 3**). Successful responses were better described by a nonlinear relationship with age, increasing through development and declining in older individuals (**Fig. 3a**; **Table S1a**). This decline coincided with a progressive increase in premature responding across the lifespan, indicating that the late-life decline in success is largely attributable to rising premature responding. Error rates remained consistently low and showed no clear age-related trajectory, whereas omissions followed a nonlinear trajectory across the lifespan, declining through early life before leveling off in older individuals. When we examined lifespan trajectories in the visual-acuity task (**Fig. S3a**), performance declined monotonically with age: success rates decreased, and errors increased, with no evidence of the nonlinear trajectory observed for sustained attention (**Fig. S4b; Table S1b)**. This contrast argues against general age-related visual decline as the primary explanation for the sustained-attention profile. In the delayed matching-to-sample task (**Fig. S3b**), however, success and premature responding followed pronounced nonlinear trajectories **(Fig. S4c; Table S1c)**. The convergent late-life increase in premature responses across both tasks suggests that age-related difficulty maintaining attentional control until target availability is preserved across distinct task contexts.

**Figure 3.**
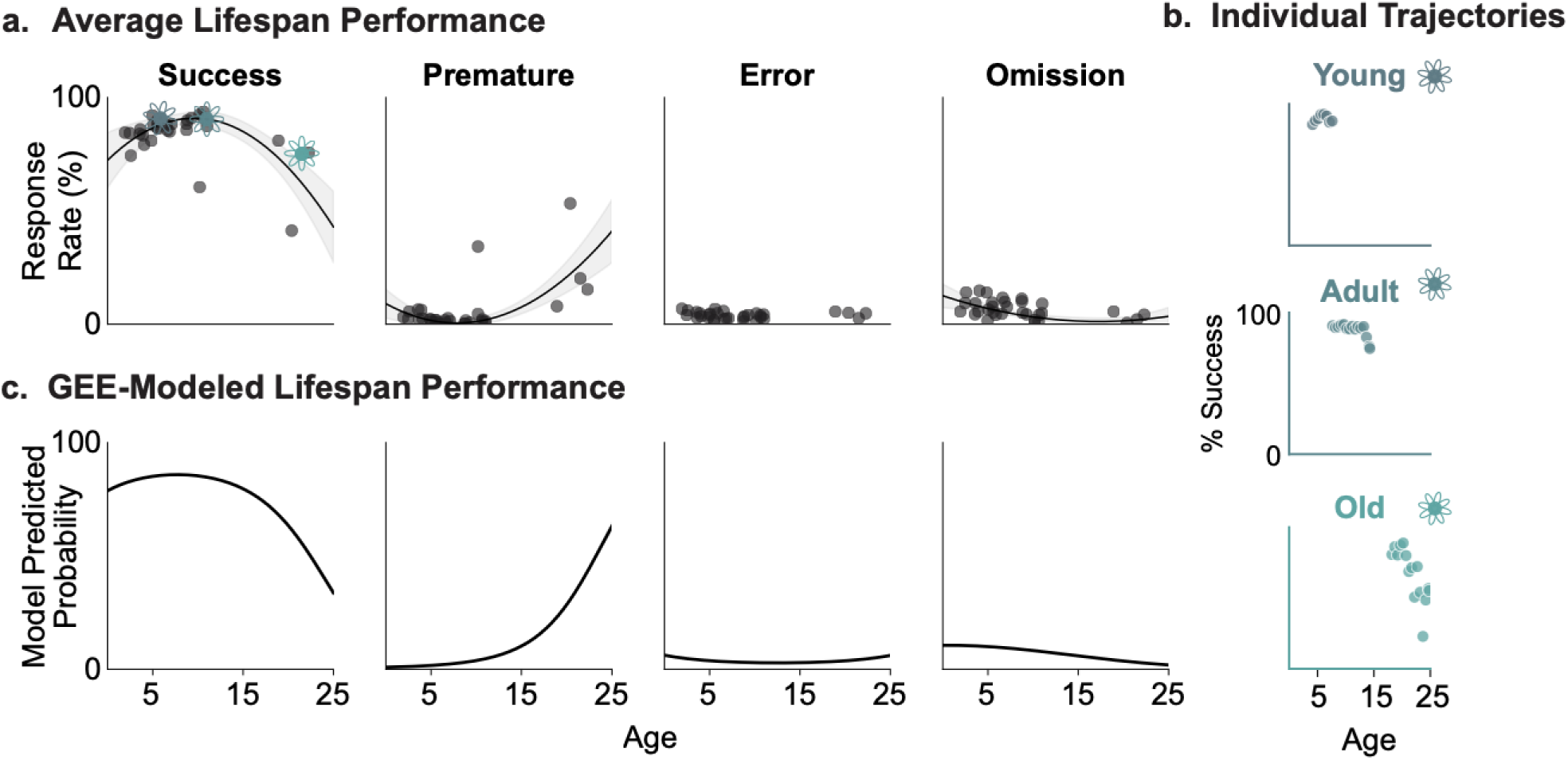
Lifespan Trajectories of Sustained Attention. **(a)** Each panel shows individual animal performance estimates (dots) plotted against age at testing (years). Regression lines with 95% confidence intervals are shown only for the selected model. When both linear and quadratic models showed a significant age effect, the quadratic model was retained only if it provided a significantly better fit than the linear model; otherwise, the linear model was retained (see Table S1 and Methods) **(b)** Representative longitudinal testing histories from young, adult, and older individuals. **(c)** Age-related predicted probabilities estimated from trial-level Generalized Estimating Equation models that account for repeated observations within individuals.

These initial analyses, however, collapsed each animal’s testing history into a single average, obscuring longitudinal changes in performance as individuals aged (**Fig. 3b**). As a result, they could not determine how these within-individual changes contributed to population-level lifespan patterns. We therefore leveraged the full longitudinal dataset, modeling 641,607 trial outcomes from 32 individuals as a function of age using generalized estimating equations and accounting for repeated observations within animals (see Methods). The longitudinal analysis recovered largely the same structure: successes, premature responses, and omissions followed quadratic age trajectories, and errors, non-significant at the population level, also followed a quadratic trajectory **(Fig. 3c**; **Table S2)**.

Together, these analyses demonstrate that sustained attention follows a robust nonlinear lifespan trajectory, evident both in animal-level summaries and in trial-level models accounting for repeated observations. This trajectory contrasted with the monotonic decline in visual acuity. The late-life increase in premature responding was also evident in an independent cognitive task, suggesting that age-related changes in the ability to maintain task-appropriate control may generalize across distinct task contexts.

### Social Status Shapes Attentional Performance, with Stronger Effects Early in Life

Beyond stable individual differences and lifespan-related change, attentional performance may also be shaped by social context. In socially structured primate societies, hierarchical status influences access to resources, social interactions, and daily experiences, making it a potentially important determinant of cognitive performance throughout life. To examine this possibility, we again leveraged the full longitudinal dataset, modeling trial-level outcomes as a function of age and social status using generalized estimating equations (see Methods).

These models estimated the independent effects of age and social status on performance, as well as their interaction, allowing us to determine whether the influence of social hierarchy varied across the lifespan.

As expected, the models confirmed significant effects of age (all main effects of age, p < 0.001**; Table S3**; consistent with **Fig. 3**). In addition, higher social status was independently associated with greater success and fewer premature responses and errors (all main effects of social status, p < 0.001; **Table S3**). Critically, these effects were qualified by significant age-by-social status interactions (**Fig. 4a**; **Table S3**). The positive association between social status and successful performance progressively weakened with age (**Fig. 4a**, left panel). Similarly, the protective effect of high social status on premature responses and errors (**Fig. 4a**, middle panels) was strongest in young individuals but progressively diminished across the lifespan, becoming negligible or even reversing among the oldest animals. Thus, the cognitive advantages associated with high social status were most pronounced early in life. In contrast, omissions exhibited a crossover interaction: higher social status predicted more omissions in younger individuals but fewer omissions in older individuals, resulting in a reversal of the status–omission relationship across the lifespan (**Fig. 4a**, right panel).

**Figure 4.**
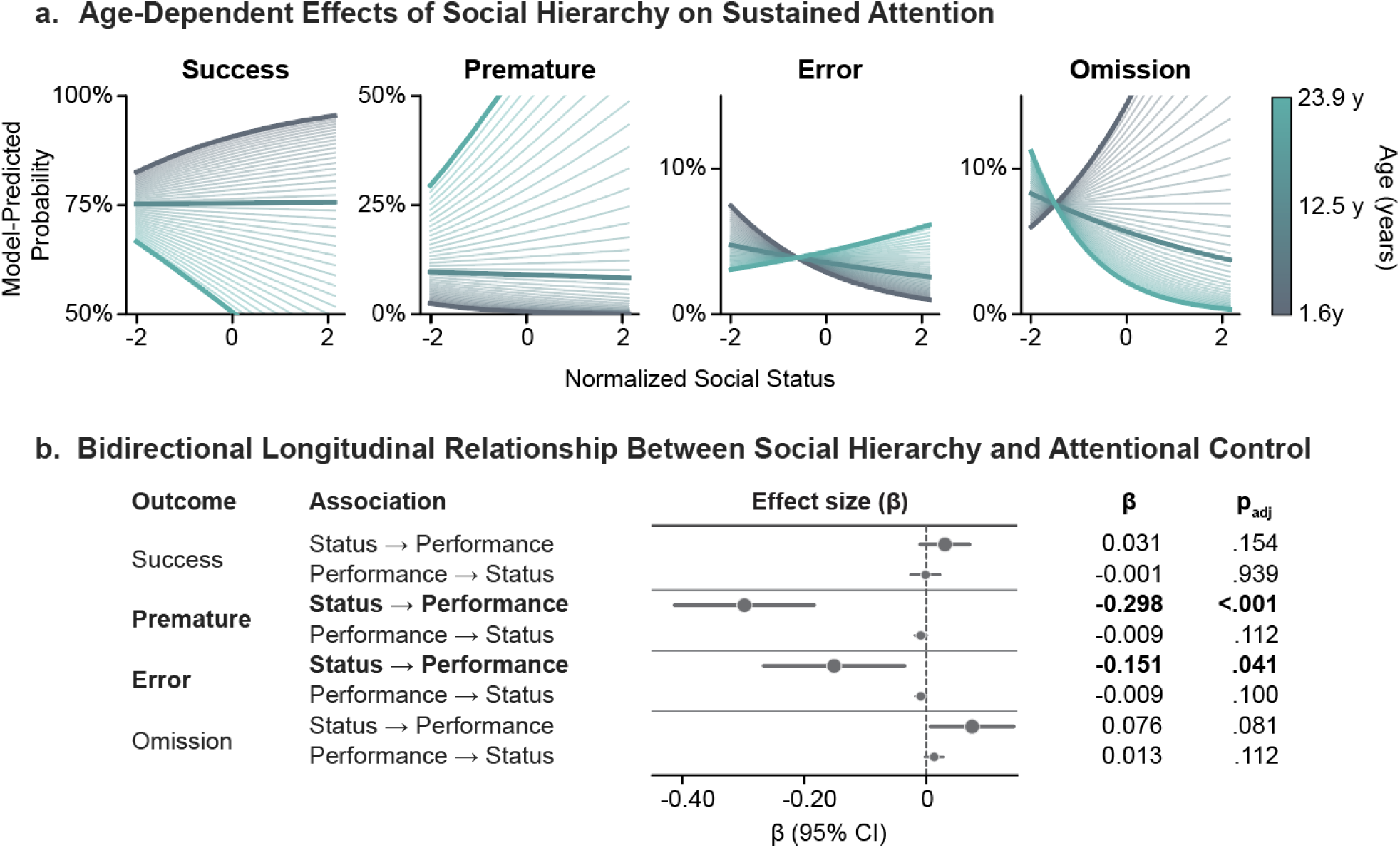
Social Hierarchy Shapes Attentional Performance Across the Lifespan. **(a)** Model-predicted probabilities from generalized estimating equation (GEE) logistic regression models including normalized social status (Elo z-score), age, and their interaction (641,607 trials; 32 M. tonkeana). Thin lines show predictions across the observed trial-level age range (1.5-25.3 years); highlighted lines correspond to representative young (1.6 y), middle-aged (12.5 y), and old (22.9 y) individuals. **(b)** Cross-lagged analyses with a 1-month lag show within-individual bidirectional associations between monthly fluctuations in social status and attentional performance. Points represent standardized β coefficients and horizontal bars denote 95% confidence intervals. Bold values indicate effects surviving FDR correction.

To examine the temporal directionality of the relationship between social hierarchy and attentional control, we tested whether month-to-month fluctuations in social status predicted future performance and vice versa (**Fig 4b**, see Methods). In brief, we summarized social status and behavioral performance at the monthly level for each animal and tested whether deviations from an individual’s typical status predicted deviations from its typical performance one month later, and conversely whether performance predicted subsequent changes in status. Higher social status prospectively predicted fewer errors (p_adj_ = 0.041) and fewer premature responses (p_adj_ < 0.001) one month later. By contrast, no measures of performance significantly predicted subsequent changes in social status after correction for multiple comparisons. The prospective effects of social status on performance were consistent across lags of one to three months. No directional effects were detected for success or omissions after correction for multiple comparisons. Together, these prospective analyses suggest that changes in social hierarchy precede changes in the response-control component of sustained attention.

Overall, these findings identify social hierarchy as a significant source of variation in sustained attention among semi-free-ranging macaques. The influence of social status was strongest early in life and was particularly evident in measures of sustained attention and premature responding. The absence of reciprocal effects from performance to subsequent social status further supports the view that social hierarchy may shape attentional control, rather than simply covarying with it. Together, these findings suggest that social hierarchy contributes to shaping attentional control throughout the lifespan of primate societies.

### Frontoparietal Connectivity Shows Lifespan Reorganization

Having established that sustained attention varies across individuals, changes across the lifespan, and is modulated by social status, we next examined whether similar principles are reflected in intrinsic brain organization. Specifically, we leveraged the group of 18 animals that underwent resting-state fMRI to test whether intrinsic functional connectivity within the cortical attention network follows lifespan trajectories that parallel those observed behaviorally.

To determine whether lifespan changes in sustained attention were mirrored at the neural level, we examined intrinsic connectivity between LIP and FEF, two core nodes of the frontoparietal attention network (**Fig. 5a**)^20,29^. Applying the same modeling framework used for the behavioral analyses, we compared linear and quadratic models of age-related variation in connectivity. LIP–FEF connectivity exhibited a significant inverted-U trajectory across the lifespan (**Fig. 5b**; **Table S4**), with the quadratic model fitting significantly better than the linear model.

**Figure 5.**
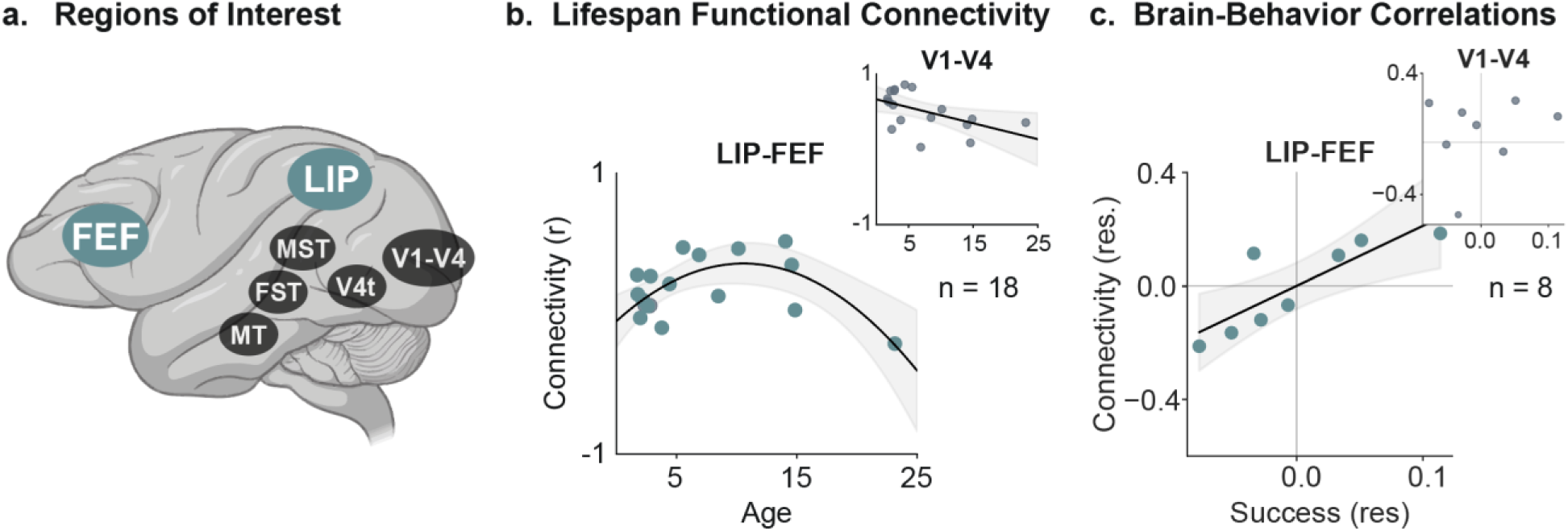
Lifespan Trajectories and Brain–Behavior Correlations in Frontoparietal Network. **(a)** Regions of interest. **(b)** Lifespan functional connectivity of the attention network. Fisher z-transformed and plotted as a function of age at scan. Main panel shows LIP–FEF connectivity; inset shows V1–V4 connectivity. **(c)** Brain–behavior relationships shown as partial correlations between connectivity and task performance after regressing both variables on age, isolating shared variance independent of development. Main panel shows LIP–FEF connectivity versus sustained attention success; inset shows V1–V4 connectivity versus sustained attention success. Regression lines and shaded 95% confidence intervals are shown only where the selected model or correlation reached statistical significance. FEF: frontal eye field; LIP: lateral intraparietal area; V1: primary visual cortex; V4: visual area 4.

A systematic examination of all measured visual and attentional connections revealed that most were either stable across the lifespan or followed monotonic linear trajectories (**Tables S4-S5**), as exemplified by V1–V4 connectivity (**Fig. 5b inset**). In contrast, the frontoparietal connection linking LIP and FEF exhibited an inverted-U profile that paralleled the behavioral lifespan trajectory.

Given the contribution of premature responding to age-related behavioral variation and the established involvement of cortico-striato-thalamic circuits in response inhibition, we additionally explored whether connectivity within this network*—*including the orbitofrontal cortex (OFC)^34^, dorsal striatum^35,36^, nucleus accumbens (NAcc)^37^, mediodorsal thalamus (MD)^34^, and centromedian-parafascicular thalamic complex (CMnPF)^38^ (**Fig. S6a***)—*showed corresponding lifespan changes. In contrast to LIP–FEF connectivity, nine of the ten examined cortico-striato-thalamic connections showed no significant linear or quadratic association with age (**Fig. S6b; Table S6**). Only DS–CMnPF connectivity exhibited a significant nonlinear trajectory (**Fig. S6b; Table S6**), providing limited evidence for systematic lifespan reorganization within this targeted circuit.

Together, these findings reveal system-specific lifespan dynamics: LIP–FEF frontoparietal connectivity exhibited an inverted-U trajectory paralleling that of sustained attention, whereas the other examined connections showed little evidence of comparable nonlinear reorganization.

### Intrinsic Connectivity Is Associated with Behavioral Performance

Finally, we asked whether variation in intrinsic connectivity is associated with individual differences in task performance. To address this question, we examined the subset of animals that underwent resting-state fMRI and completed the attention task within six months before scanning (n = 8; **Fig. S7**).

Consistent with its proposed role in sustained attention, stronger LIP–FEF connectivity was associated with higher success rates (**Fig. 5c**). Because both behavior and connectivity varied with age, we repeated the analysis using partial correlations controlling for age. The association between LIP–FEF connectivity and success remained significant, whereas no relationships were observed for errors or premature responses. Importantly, neither connectivity within the control visual pathway (V1–V4) nor any visuo-visual connection was associated with success rates after controlling for age (**Fig. 5c inset**), suggesting that the LIP–FEF association did not simply reflect global variation in functional connectivity.

We next broadened our analysis beyond the LIP–FEF and V1–V4 comparisons to examine all measured functional connections within the frontoparietal attention and visual systems (**Table S7**). Given the limited sample size and exploratory nature of these broader analyses, p-values are reported uncorrected for multiple comparisons. Exploratory associations showed some anatomical organization: success rates were preferentially associated with FEF-centered connectivity, omissions with LIP-centered connections, and errors with connections among visual regions. Although these observations require replication in larger samples, their apparent anatomical organization suggests that distinct components of task performance may rely on partially dissociable functional systems. Finally, to complement the exploratory characterization of brain–behavior relationships, we examined cortico-striato-thalamic connectivity (**Table S8**). Within this network, the clearest associations involved OFC-centered connections: stronger OFC–CMnPF connectivity was associated with fewer omissions, the only association to survive FDR correction, whereas stronger OFC–NAcc connectivity was associated with fewer premature responses. Additional associations involved OFC–MD connectivity with premature responses and NAcc–dorsal striatum connectivity with omissions (**Table S8**). Together, these exploratory findings highlight specific cortico-striato-thalamic connections for targeted investigation in larger samples.

Together, these findings suggest that brain–behavior associations were not uniformly distributed across the examined connections. Most notably, frontoparietal connectivity was associated with successful sustained-attention performance, whereas visual connectivity was more often linked to target-selection errors. Although preliminary, the broader pattern suggests that distinct components of task performance may relate to partially dissociable aspects of intrinsic functional organization.

## Discussion

The present study offers an integrated account of sustained attention across multiple scales of biological organization. By combining long-term cognitive phenotyping in semi-free-ranging macaques with measures of social hierarchy and intrinsic brain connectivity, we examined how stable individual differences, developmental trajectories, social environments, and neural circuits jointly shape attentional function within the same population. Rather than reflecting isolated levels of explanation, our findings indicate that these dimensions form a coherent organizational framework through which sustained attention emerges and adapts over time. More broadly, they illustrate how longitudinal studies in naturalistic primate societies can uncover principles of cognitive organization that are difficult to resolve in either traditional laboratory models or cross-sectional human studies.

Within this multiscale framework, sustained attention was jointly organized by stable individual phenotypes and conserved developmental trajectories. While most animals consistently exhibited high levels of attentional control over several years, a subset persistently displayed elevated premature responding, indicating that the tendency to respond before target onset can represent a stable behavioral trait rather than a transient state. The stability of these behavioral profiles across years, together with their generalization across cognitive tasks, indicates that sustained attention contains a trait-like component that persists despite ongoing behavioral and contextual fluctuations. This pattern closely parallels human evidence showing that impulsive-response tendencies can represent a stable individual characteristic over time, displaying high test–retest reliability^39^, and that attention-control abilities generalize across tasks^5^. In humans, persistent inter-individual differences in attentional and response control span a continuum from normative variation to neurodevelopmental disorders such as ADHD^40^, which often persists from childhood into adulthood^41,42^. Our findings demonstrate that similarly stable attentional phenotypes also emerge within an otherwise healthy macaque population, suggesting that stable inter-individual variation in attentional control may be an intrinsic feature of primate cognition rather than exclusively a hallmark of pathology. More broadly, these findings support the view that stable individual differences are not merely residual variability around a common cognitive architecture but constitute a fundamental dimension of cognition^43,44^.

Complementing these stable individual phenotypes, sustained attention also exhibited a characteristic developmental organization across the lifespan. Attentional performance followed a characteristic inverted-U trajectory, improving from childhood through adolescence, peaking in adulthood, and declining in old age. Importantly, these developmental changes unfolded alongside long-lasting individual behavioral profiles, suggesting that maturation modifies attentional capacity while stable inter-individual differences remain an important source of variation. Within this lifespan trajectory, premature responding increased with age, consistent with reports of age-related impairments in response inhibition in rhesus macaques performing analogous attention tasks^45^. The parallel increase in premature responding in the delayed matching-to-sample task further suggests that this age-related change generalizes across cognitive contexts. This lifespan profile extends previous laboratory studies, which have primarily compared isolated developmental stages^45,46^, by reconstructing a continuous lifespan trajectory within a single semi-free-ranging population tested under a common behavioral paradigm. As in natural populations, age classes were not evenly represented, with relatively fewer young and old individuals. Although this uneven sampling may reduce the precision of the estimated trajectory at the extremes of the lifespan, it preserves the naturally occurring demographic composition of an existing macaque social group, including the progressive attrition of individuals at older ages^47,48^, in contrast to most human lifespan studies, which typically recruit balanced age cohorts by design (e.g., Cam-CAN; HCP-Aging)^49,50^. This highlights an important trade-off in lifespan research: whereas experimentally balanced cohorts provide greater precision for estimating age-related trajectories, naturalistic populations offer a more ecologically representative picture of how cognitive functions are distributed across the lifespan. Notably, peak attentional performance coincided with the prime reproductive period in macaques^51,52^, when individuals may face substantial demands on attentional resources for mate competition, offspring care, and maintaining their social position. Although speculative, this temporal correspondence raises the intriguing possibility that sustained attention may be tuned to the changing ecological pressures encountered across the lifespan. Perhaps most interestingly, the lifespan trajectory broadly mirrors that observed in humans, in whom sustained attention continues to improve through young adulthood and peaks in the early forties^4^, despite profound differences in ecology, environmental experience, and life history between the two species. Rather than reflecting species-specific developmental patterns, this cross-species convergence is consistent with the existence of evolutionarily conserved developmental programs underlying the maturation and aging of sustained attention.

Beyond stable individual differences and developmental change, our findings identify social status as an additional level of organization shaping sustained attention. Young higher-ranking individuals consistently exhibited superior attentional performance and reduced impulsive responding, although these advantages were strongest early in life, indicating that attentional capacities are embedded within the social environment in which they develop and operate. Importantly, longitudinal analyses showed that higher social status prospectively predicted fewer premature responses and fewer errors one month later, suggesting that social hierarchy can exert a lasting influence on attentional control. These findings extend a broader primatological literature showing that vigilance in free-ranging macaques varies with dominance rank and genotype^53^, is modulated by kin availability^54^, and can spread through social relationships via vigilance contagion^55^. One possible explanation is that social subordination is associated with chronic stress^56,57^ and altered prefrontal function^58^, which could impair the cognitive processes underlying task performance. Although the underlying biological pathways remain to be established, this behavioral pattern closely parallels findings from early life in humans, where higher socioeconomic status predicts better attentional and executive functioning^6,59^, and fewer impulsive and attention-related problems^6,60^. Importantly, in humans, lower performance in nonsocial cognitive tasks does not necessarily imply poorer cognitive capacity: social status may instead shape which sources of information are prioritized according to their relevance in an individual’s social environment^61^. From this perspective, the attentional differences observed here could partly reflect differences in the allocation of attentional resources across positions within the social hierarchy. Together, these findings suggest that the influence of social status on sustained attention may represent a general feature of primate cognition, despite marked differences in the organization of human and macaque societies. This social organization of attention is difficult to capture in conventional laboratory studies, where animals typically live in small, experimentally constrained groups with limited opportunities for natural social dynamics. Likewise, human studies rarely permit repeated longitudinal assessment of cognitive performance together with objectively quantified social hierarchy within stable societies. By combining long-term behavioral phenotyping with natural social organization, our study reveals that sustained attention is shaped not only by intrinsic biological factors but also by the social environment in which cognition is expressed.

The behavioral organization of sustained attention was mirrored by the organization of its underlying frontoparietal network. At the population level, frontoparietal functional connectivity followed a similar nonlinear developmental trajectory as attentional performance, peaking in adulthood and declining with age. Notably, the connectivity peak occurred slightly later than the behavioral peak, potentially reflecting continued maturation and consolidation of the frontoparietal network beyond the period of maximal behavioral performance. Beyond these developmental effects, LIP–FEF connectivity also captured inter-individual differences in sustained attention. This relationship persisted after controlling for age, demonstrating that variation in frontoparietal network strength is associated with attentional performance independently of developmental stage. Importantly, both the nonlinear lifespan trajectory and the relationship between LIP–FEF connectivity and attentional performance were most clearly observed in frontoparietal connectivity, with limited evidence for comparable effects in purely visual networks, indicating that these effects cannot be explained by generalized changes in sensory processing. Although the brain–behavior analyses were based on a limited imaging sample and should therefore be interpreted cautiously pending replication, the anatomical specificity of these associations argues against a nonspecific effect of global functional connectivity. Consistent with this specificity, selected cortico-striato-thalamic connections showed comparatively limited lifespan reorganization despite the age-related increase in premature responding. Nevertheless, exploratory brain–behavior associations between OFC-centered connectivity and premature responding suggest that individual variation within this circuitry may contribute to response control even in the absence of systematic age-related reorganization. Thus, while cortico-striato-thalamic connectivity may contribute to individual differences in response control, the frontoparietal network uniquely captured both the developmental trajectory of sustained attention and inter-individual differences in attentional performance. These findings extend a large body of causal, lesion, neurophysiological, and neuroimaging evidence implicating frontoparietal networks in attentional control^26,28,29,62–66^ by demonstrating that the intrinsic organization of this canonical attention network also captures variation across both the lifespan and individuals. This finding parallels human studies showing that distributed patterns of functional connectivity are associated with individual differences in sustained attention across both adult and developmental populations^67–69^. Age-related changes in this network organization may provide a neural basis for the behavioral decline observed later in life: in humans, aging is associated with reduced functional connectivity within frontoparietal control networks and poorer attentional control^70^. Converging evidence from macaques further shows that aging alters prefrontal cortical function and synaptic organization^71,72^, providing a potential biological substrate for the decline in frontoparietal connectivity observed here. Together with comparative evidence for a broadly conserved organization of attention networks across primates^25^, alongside species-specific elaborations^73^, our findings identify an evolutionarily conserved neural architecture underlying the multiscale organization of sustained attention.

Overall, our findings support a view of sustained attention as a multiscale biological function rather than a unitary cognitive process. Stable individual differences, developmental trajectories, social status, and intrinsic brain networks each contribute distinct yet complementary levels of organization that together shape attentional behavior. By combining long-term cognitive phenotyping in semi-free-ranging macaques with functional neuroimaging, this study complements emerging collaborative efforts to build large-scale, shared non-human primate resources^74,75^, and demonstrates how comparative, integrative, naturalistic approaches can bridge levels of analysis that remain difficult to integrate in either conventional laboratory models or cross-sectional human studies. In doing so, our findings establish sustained attention as a function organized across biological scales, from stable individual variation and lifespan development to social structure and intrinsic brain organization.

## Acknowledgements

The authors are grateful to the University of Strasbourg, the CNRS, and SILABE (silabe.com) for their support of this research and for providing expert animal care, especially Pierrick Regnard for veterinary support around MRI acquisitions. This work was funded by the Swiss National Science Foundation through an SNSF Starting Grant to I.S. (grant no. 211431) and by ANR-21-CE37-0016 to S.B. and H.M., and was co-funded by the French State-Region contract CPER I2MT (2014–2021), R-IRM (2021–2027), and the European Union through the European Regional Development Fund (FEDER Grand Est). MRI scans were performed on the IRIS platform of the ICube lab, a member of the France Life Imaging network (grant ANR-11-INBS-0006). We sincerely thank Julien Lamy, Paulo Loureiro de Sousa, and Aurore de Cauwer for their invaluable assistance during every stage of MRI data acquisition, and Marion Sourty for her expert sanity checks of the fMRI data. The development of the MALT was supported by the University of Strasbourg Institute for Advanced Study (USIAS) as part of a USIAS fellowship to H.M. We also extend our thanks to Adam Rimelé for his computer and programming support for the MALT, to Jonas Fizet for his contribution to MALT engineering, and to Quentin Fuchs for his preliminary analysis of the MALT data in relation to socio-demographic factors. We also thank Michael Schmid for his comments on an earlier version of the manuscript.

The authors declare no competing interests.

## Ethics Statement

NHPs were monitored in accordance with the guidelines of the European Directive 2010/63 and handled in strict accordance with good animal practice as defined by the French National Charter on the Ethics of Animal Experimentation. All animal protocols were approved by the French Ministry of Higher Education and Research (permit number: APAFIS 2023062617104978_v3) and by the Primate Centre ethics committee (registration n°B6732636) in compliance with EU Directive 2010/63/EU.

## Materials and Methods

### Subjects and Study Site

Subjects comprised 32 *Macaca tonkeana* (12 females) living in semi-free-ranging conditions and spanning juvenile, adult, and aged individuals. Animals were tested repeatedly over several years, so age is indexed as the midpoint of each individual’s testing history in animal-level analyses (1.9–22.3 years) and as age on the day of the trial in trial-level analyses (1.5–25.3 years). Because wild and captive macaques typically live up to 25–30 years, with the maximum recorded longevity for *M. tonkeana* in captivity reported at 28.4 years^13^, the sampled age range spans most of the species’ natural lifespan. Animals were group-housed at the Primate Centre of the University of Strasbourg – Simian Laboratory Europe (SILABE), in a 3,788 m² wooded park with indoor-outdoor shelters. All behavioral observations were collected based on voluntary participation of the animals during a period of up to 7 years (**Fig. S1**), and the MRI procedure was approved by the Primate Centre ethics committee (registration n°B6732636) in compliance with EU Directive 2010/63/EU.

### Behavioral Data Collection

#### Apparatus

Behavioral data were acquired using Machines for Automated Learning and Testing (MALT), custom-built touchscreen devices developed in-house at the Primate Centre of the University of Strasbourg, and integrated into the group’s naturalistic enclosure (**Fig. 1a-b**)^30,31^. Each unit integrates a touchscreen interface, a fluid reward system, a dual Radio-Frequency Identification (RFID) system for automated behavior logging^30^, and a video camera. The apparatus enabled spontaneous, unsupervised task participation under naturalistic conditions with minimal experimenter interference^30,31^. The animals had access to up to four touchscreen units. All devices were accessible *ad libitum* throughout the 24-hour period (except during cleaning procedures twice per week), enabling voluntary, *in situ* task participation. Each device featured tasks with graded difficulty levels varying in stimulus parameters (duration, luminosity, delay). In the current paper, data were analyzed only after training phases, when performance stabilized at advanced levels, ensuring measurements reflected established behavior rather than learning.

#### Cognitive Tasks

To assess sustained attention, we ran a touchscreen-based adaptation of the 5-CSRT task, a cross-species analog of the human CPT (**Fig. 1c**)^32,33^. Each trial began with the animal touching a yellow square. After 500 ms, five white hollow circles appeared in a circular configuration on a black background. Following a variable delay (1500–3000 ms), a yellow dot was flashed for 25 ms at two luminance levels (100% or 60%). Premature touches during the delay triggered a 4000 ms white-screen timeout. After the flash, the animal had 5000 ms to touch the target location. Correct responses were rewarded with 1 mL of diluted syrup (1:10) delivered over 2000 ms; incorrect responses resulted in a 4000 ms white-screen timeout. All responses were followed by 500 ms of auditory feedback.

Two additional control tasks were administered. Reward and timeout contingencies matched those used in the 5-CSRTT. The first control task was a visual acuity task assessing basic visual discrimination using reduced-size stimuli (**Fig. S3a**). After touching a green start square and a 500 ms delay, a target stimulus was randomly selected from a pool of 7 shapes and presented at reduced size. Upon touching the target, three shapes were presented while the target remained visible. Animals had 5000 ms to select the target-matching shape. Premature touches during the delay period (i.e., before the target stimulus appeared) triggered a timeout. The second control task was a delayed object matching task requiring short-term maintenance of object identity and subsequent attentional selection among distractors^76^ (**Fig. S3b**). Each trial began with touching an orange start square, followed by a 500 ms delay. A target shape (randomly selected from a set of 4 shapes) appeared for 3000 ms (sample phase). Following a 1000 ms delay, all four shapes reappeared, and the animal had 5000 ms to select the original target. Premature touches during the sample phase triggered a timeout. Across all tasks, behavioral performance resulted in four possible outcomes (**Fig. 1d**): *successes*, correct selections of the target stimulus; *premature responses*, touches made before the response phase began; *errors*, selections of non-target stimuli; and *omissions*, when no response was made within the allotted window. Performance was assessed in terms of accuracy. Responses initiated within 100 ms of trial onset were reclassified as premature, as such latencies fall below the threshold for stimulus-driven cognitive processing^77^.

Animals voluntarily engaged with touchscreen tasks over multiple years (December 2016-January 2024), producing thousands of self-initiated sessions (54–7’973; mean = 2’537) and attempts (837–73,064; mean = 20,041) per individual, with a median of 4 attempts per session (IQR: 2–8; see also **Fig. S1**). Three individuals were excluded from analyses: two due to an RFID identification error that assigned the same chip ID to both animals, making their behavioral data indistinguishable, and a third for never having engaged with the primary task.

### Quantification of Social Status

We quantified social status using a semi-automated Elo-rating system based on touchscreen displacement events, previously validated against direct behavioral observations in *M. tonkeana*^31^ and in *Papio papio*^78^. Specifically, a displacement event was recorded whenever one individual relinquished access to the touchscreen following the approach of another individual, allowing the latter to occupy the device. These displacement interactions were treated as win–loss outcomes and automatically logged via subcutaneous RFID microchips. Elo scores were computed using a standard algorithm, with all individuals initialized at 1000; following each interaction, the displacing individual gained points and the displaced individual lost points^31,79^.

### Behavior Analysis

We analyzed performance as the proportion of each response type per session: successes, premature responses, omissions, and errors. These metrics served as the primary outcome measures for three behavioral analyses targeting, respectively, individual behavioral phenotypes, lifespan trajectories, and the effects of social status on task performance.

First, to identify distinct individual behavioral phenotypes, we visualized longitudinal trajectories of task performance for each subject using stacked proportion plots displaying the distribution of the four response types across consecutive testing sessions, revealing stable individual differences in attentional capacity and impulsivity that persisted across time. To formally test the stability of these phenotypes, we computed success and premature response rates within non-overlapping 6-month windows spanning each animal’s testing history, including individuals with at least 4 such windows (n = 26), and calculated the intraclass correlation (ICC) across these windows^80^.

Second, to characterize lifespan trajectories, we modeled behavioral performance as a function of age using weighted least squares linear and quadratic regression models, with each data point representing a single animal’s mean performance across valid trials, weighted by each animal’s number of attempts.

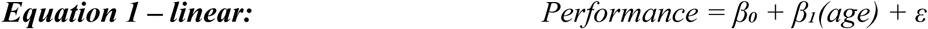

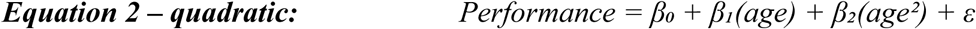

where *Performance* is the mean behavioral measure for each animal, *age* is the animal’s age in years, calculated as the temporal midpoint between each individual’s first and last trial (1.9-22.3 years), *β₀* is the intercept, *β₁* is the linear effect of age, *β₂* is the quadratic effect of age, and *ε* is the residual error. Model fit was quantified using adjusted *R²*. Linear and quadratic models were first evaluated separately for a significant effect of age (*β₁* for the linear model, *β₂* for the quadratic model; α = 0.05). If only one model showed a significant age effect, that model was retained. If both models were significant, they were compared using a nested F-test, and the quadratic model was retained only if it provided a significantly better fit than the linear model; otherwise, the linear model was retained as the more parsimonious solution. If neither was significant, no model was retained. Regression lines are displayed only for significant models. Full model statistics are reported in **Table S1.**

Third, to examine age-related and social-status-related effects on task performance at the trial level, we used Generalized Estimating Equations (GEE). GEE extends standard regression to repeated-measures data by accounting for the non-independence of repeated trials from the same individual. Trials were clustered by individual and testing day, with an exchangeable working correlation structure and a binomial link function. Because age and social status both varied over the course of the study—age increasing continuously, Elo ratings fluctuating as the hierarchy changed—each trial was assigned the individual’s age and Elo rating for the calendar day on which it occurred (observed trial-level age range: 1.5-25.3 years). Thirty-five trials fell on days for which no Elo rating was not available for that individual and were excluded, leaving 641,607 trials from 32 individuals.

We first fitted a GEE model including linear and quadratic age terms (Equation 3). This model builds on the age-based population-level analyses described above while accounting for the repeated-measures structure of the data, allowing us to test whether the curvature identified above replicated at the trial level.

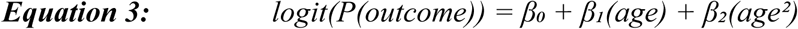

where P(outcome) is the probability of a given behavioral outcome (success, premature response, error, or omission), β₀ is the intercept, β₁ is the linear effect of age, and β₂ is the quadratic effect of age. Age was mean-centered prior to squaring.

To test whether social status shaped performance, and whether its influence varied across the lifespan, we fitted a second GEE model with age, standardized social status (Elo rating), and their interaction:

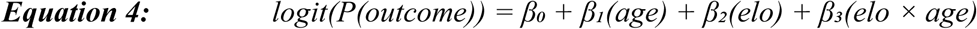

where *P(outcome)* is the probability of a given behavioral outcome, *β_0_* is the intercept, *β_1_* and *β₂* estimate the main effects of age and social status, respectively, and *β_3_* estimates their interaction. Elo ratings were standardized (mean = 0, SD = 1). Age was mean-centered, such that *β₂* reflects the effect of social status at the average age of the sample.

The Benjamini–Hochberg false discovery rate (FDR; α = 0.05) correction was applied separately within each model: across the 8 coefficients of the age model (linear and quadratic age for each of the four behavioral outcomes) and across the 12 coefficients of the social status model (age, social status, and their interaction for each outcome); all reported effects survived correction^81^. Full model statistics are reported in **Tables S2-S3**.

To examine whether changes in social status preceded changes in performance, or vice versa, we first summarized the data at the monthly level. For each animal and each calendar month, we calculated average social status (Elo-rating) and average task performance. We then asked whether months in which an animal occupied a higher (or lower) social rank than usual were followed by months in which it performed better (or worse) than usual, and conversely, whether months with better or worse performance were followed by changes in social status. To isolate these within-individual fluctuations, both social status and behavioral measures were expressed relative to each animal’s own average across the study period. Thus, the analyses tested whether deviations from an individual’s typical status predicted subsequent deviations from its typical performance, rather than whether high-ranking animals generally differed from low-ranking animals. These lagged relationships were estimated using cross-lagged regression models^82^ with a one-month delay between predictor and outcome, including the previous month’s value of the outcome and age as covariates. P-values were corrected across all eight tests using the Benjamini–Hochberg false discovery rate procedure (α = 0.05).

### Neuroimaging Data

#### Acquisition

Twenty-one animals underwent MRI scanning. Three subjects with frontal signal dropout persisting after TOPUP correction were excluded by visual inspection^83–85^. The final sample consisted of 18 *M. tonkeana* (11 females; aged 1.7–23.2 years at the time of scanning). Of these, n = 8 also belonged to the behavioral cohort.

Animals were sedated and monitored throughout scanning to minimize motion artifacts. Physiological monitoring included an MRI-compatible optical temperature probe (OpSens OTG-M1400), a fiber-optic pulse oximeter for SpO2 and heart rate (Nonin 7500FO-3), and continuous video surveillance. Animals were placed under assisted mechanical ventilation with parameters adjusted to body weight (tidal volume: 10–20 mL/kg; respiratory rate: 20–40 cycles/min; I/E ratio: 1:3). General anesthesia was maintained with inhaled isoflurane, which was progressively lowered from 1.5% to approximately 1% over the course of the anatomical acquisition preceding rs-fMRI, targeting the 0.8–1% range associated with lighter anesthesia depth in macaques^25^. Scans were conducted using a Siemens Magnetom Vida 3T MRI scanner with a 32-channel human head coil pre-tested on post-mortem macaque brains. Sessions lasted approximately 20 minutes and included T1-weighted, T2-weighted, and resting-state functional MRI (rs-fMRI) acquisitions; the analyses presented here focus on rs-fMRI data.

Imaging parameters for rs-fMRI scans were as follows: TR/TE = 1450/50 ms, flip angle = 67°, field of view = 120 × 120 mm, voxel size = 1.5 × 1.5 × 1.5 mm isotropic, 48 slices, multiband acceleration factor = 4, 830 volumes (∼20 minutes). In the full dataset, most subjects were scanned with this protocol, except for the first nine macaques, for which acquisition was limited to 630 volumes (approximately 15 minutes) due to an earlier version of the protocol. Anatomical scans used a 3D MPRAGE sequence: TE/TR = 2.71/2400 ms, flip angle = 9°, field of view = 160 × 160 × 112 mm³, voxel size = 0.5 mm isotropic.

#### Preprocessing

Functional images were corrected for distortion using FSL^84^ *topup* and for motion using AFNI^86^ *3dvolreg*. Anatomical images were skull-stripped using the nBEST pipeline and registered to the NIMH Macaque Template (NMT) v2.1 using ANTs^87–89^. Functional data were aligned to individual anatomy using rigid and affine ANTs transformations^88^. The 4D functional timeseries were despiked using AFNI *3dDespike*^86^. Adaptive motion censoring removed approximately the worst 5% of volumes based on framewise displacement computed using a 25 mm radius to account for macaque brain dimensions^90^. Motion confounds were modeled using six rigid-body transformation parameters (three translations and three rotations), along with their temporal derivatives.

Quality control excluded subjects with mean motion exceeding 3.5 mm or >15% censored volumes. Across the 18 macaques, mean framewise displacement was 0.76 ± 0.54 mm (range 0.29–1.95 mm), and retained volumes ranged from 572 to 748. Preprocessed functional data were transformed onto the NMT space by concatenating the transformation matrices^89^. Spatial smoothing used a 2mm FWHM Gaussian kernel. Denoising combined bandpass filtering (0.01-0.1 Hz), quadratic polynomial detrending, and motion parameter regression using AFNI *3dTproject*^86^.

#### Regions of Interest

Functional connectivity was assessed using anatomically defined ROIs for two functional systems. ROI masks were derived from the Cortical Hierarchy Atlas of the Rhesus Macaque (CHARM) and the Subcortical Atlas of the Rhesus Macaque (SARM), both aligned to the NMT template^89,91^. Mean BOLD signal within each ROI was extracted to generate seed time series. Functional connectivity between regions was quantified using pairwise Pearson correlation coefficients.

Because the frontoparietal attention network primarily supports sustained spatial attention, we used functional connectivity between the lateral intraparietal area (LIP) and frontal eye fields (FEF) as our primary measure of frontoparietal network organization, as reciprocal interactions between these regions form a core circuit for spatial attention and oculomotor control^92^. As a control, we examined connectivity within the visual system, including visuo-visual pathways such as V1–V4 (**Fig. 5a**), which support the early visual processing required for our tasks, but are not centrally implicated in attentional control. This comparison allowed us to determine whether observed effects were specific to attention-related networks rather than reflecting global changes in functional connectivity. In contrast, premature responding is thought to rely on distributed cortico-striato-thalamic connections rather than a single canonical pathway **(Fig. S6a)**. We therefore examined all pairwise functional connections among five regions consistently implicated in inhibitory control—the orbitofrontal cortex (OFC)^34^, dorsal striatum^35,36^, nucleus accumbens (NAcc)^37^, mediodorsal thalamus (MD)^34^, and centromedian-parafascicular thalamic complex (CMnPF)^38^. As a quality-control step, we examined the group-average functional connectivity matrix (**Fig. S5**). As expected under anesthesia, connectivity was strongest among neighboring visual regions and weaker for long-range association connections, consistent with previous resting-state studies in non-human primates^25,93^.

#### Brain Analyses

First, to examine lifespan trajectories, we modelled Fisher z-transformed functional connectivity as a function of age using ordinary least squares linear and quadratic (inverted-U) regression models.

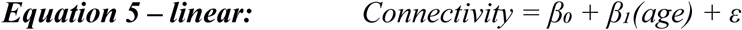

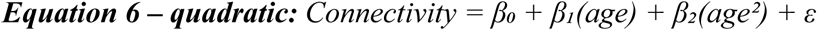

where *Connectivity* is the Fisher z-transformed functional connectivity, *age* is the animal’s age (years) at the time of MRI acquisition, *β₀*is the intercept, *β₁*is the linear effect of age, *β₂*is the quadratic effect of age, and *ε* is the residual error. Model selection followed the identical procedure used for the behavioral lifespan analyses above (adjusted *R²* for fit, α = 0.05 significance threshold for *β₁*/*β₂,* nested *F-test* when both were significant). Full model statistics are reported in **Tables S4-S6**.

Second, to examine brain-behavior relationships, we integrated behavioral data from a subset of scanned subjects (n = 8; 4 females; age range 2.8–15.3 years). Behavioral metrics were extracted within a six-month window preceding each subject’s scan date (minimum 100 trials; mean = 288 ± 219 trials per subject; mean temporal distance to scan = 98.6 ± 45.7 days; see **Fig. S7**). Four outcome measures were quantified: success, errors, omissions, and premature responses. Brain–behavior relationships were assessed using Pearson correlations between behavioral measures and connectivity. Partial correlations controlling for age were additionally calculated. Analyses were replicated for 1- and 2-year windows preceding the scan. Statistical significance was evaluated at α = 0.05 throughout.

