## Supplementary Information for "Attention Across Scales: From Individual Variation to Social Hierarchies and Brain Networks in Semi-Free-Ranging Macaques"

**Authors and affiliations**

Simona Vaitekunaite[1], Sarah Silvère[2], Hélène Meunier[2], Sébastien Ballesta[2]*, Ilaria Sani[1]*†

[1] Department of Basic Neurosciences, University of Geneva, CH

[2] LNCA (UMR 7364), University of Strasbourg, Strasbourg, FR

* co-last authors

† corresponding author

Ilaria Sani, Department of Basic Neurosciences

Campus Biotech, Chemin des Mines 9, 1202 Geneva, Switzerland

**Keywords:** Sustained attention, Attentional traits, Lifespan trajectories, Resting-state functional connectivity, Social hierarchy, Semi-free-ranging macaques; neuroethology; naturalistic behavior

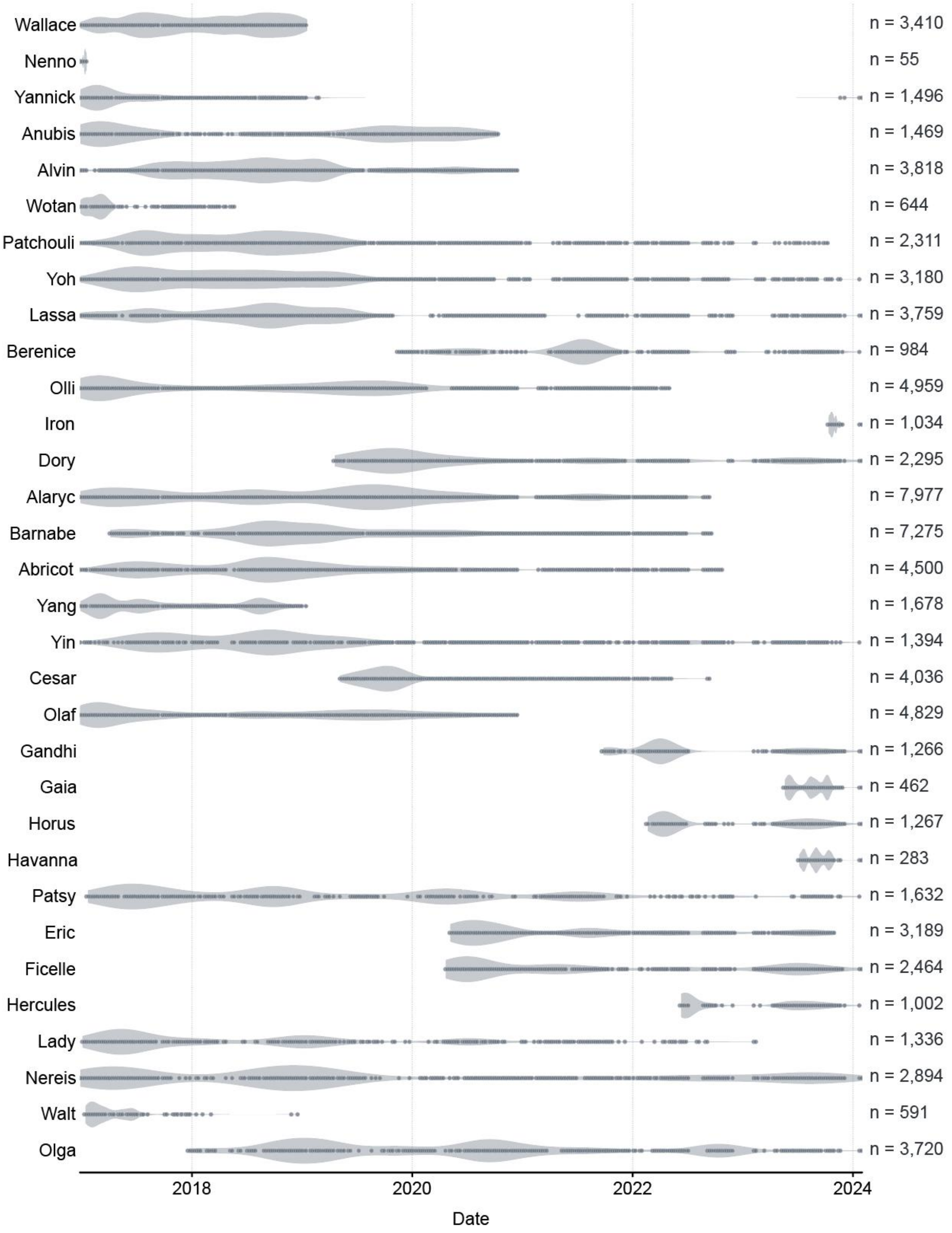


***Figure S1. Longitudinal task engagement across the M. tonkeana behavioral cohort.*** *Each row represents one individual (n = 32), restricted to the period with overlapping hierarchy (Elo) data. Rows are ordered from highest to lowest mean success rate. Markers indicate weeks containing ≥1 testing session; the shaded violin reflects testing density over time, weighted by daily attempt count. Total session count (n) for each individual is given on the right. The x-axis spans the full recording period across the cohort. A testing session represents a voluntary connection to the MALT device, during which individuals perform cognitive tasks.*

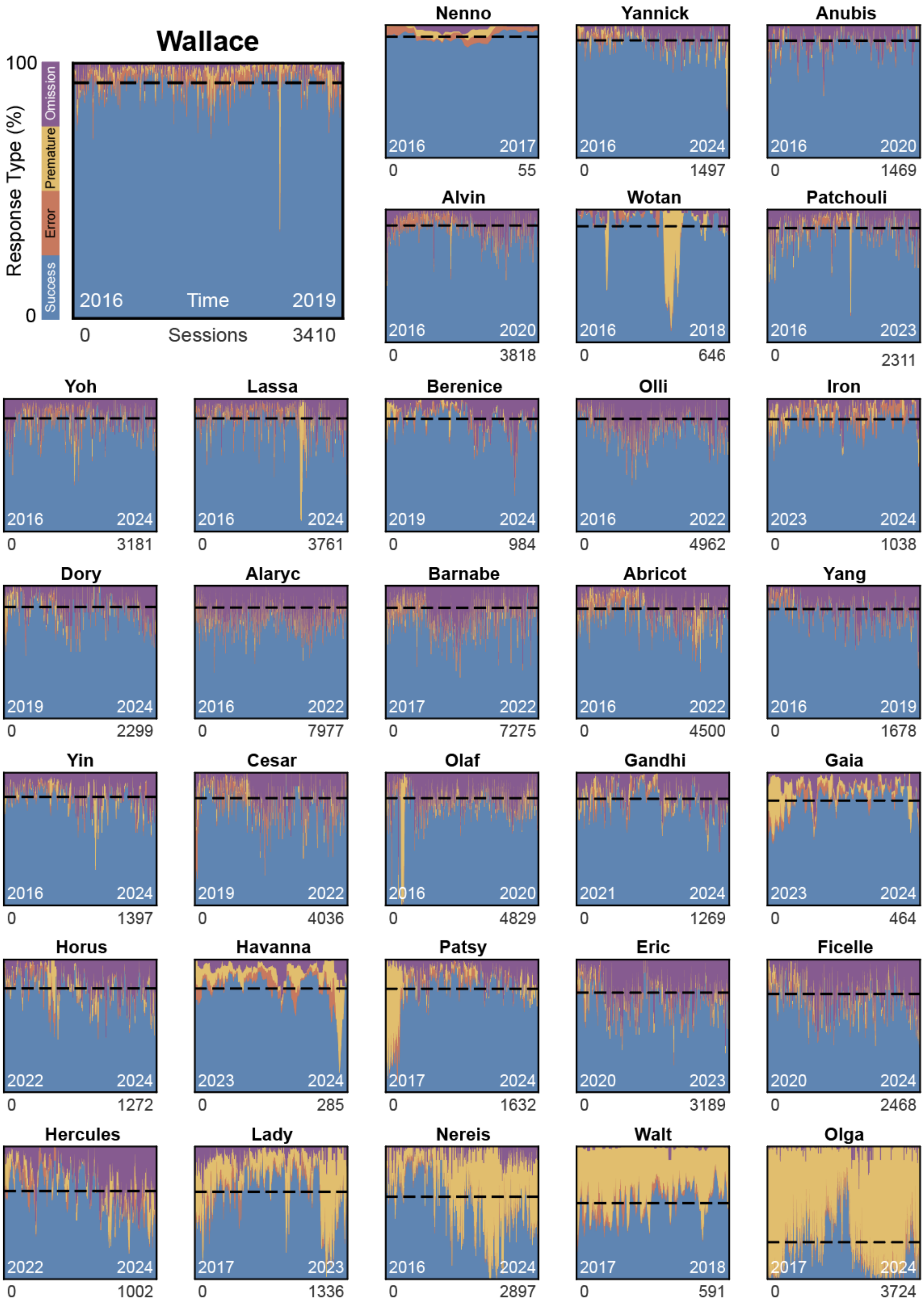


***Figure S2. Session-level behavioral trajectories for each individual.*** *Each panel shows one individual, ordered from highest to lowest mean success rate. The start and end years of data collection are shown inside each panel (bottom left/right, white), and the total number of sessions is noted below the axis (bottom right). Stacked shaded areas represent the rolling proportion of response outcomes across sessions (window = 10 sessions); colors indicate successes (blue), errors (orange), premature responses (yellow), and omissions (purple). The horizontal dashed line indicates each individual's overall mean success rate.*

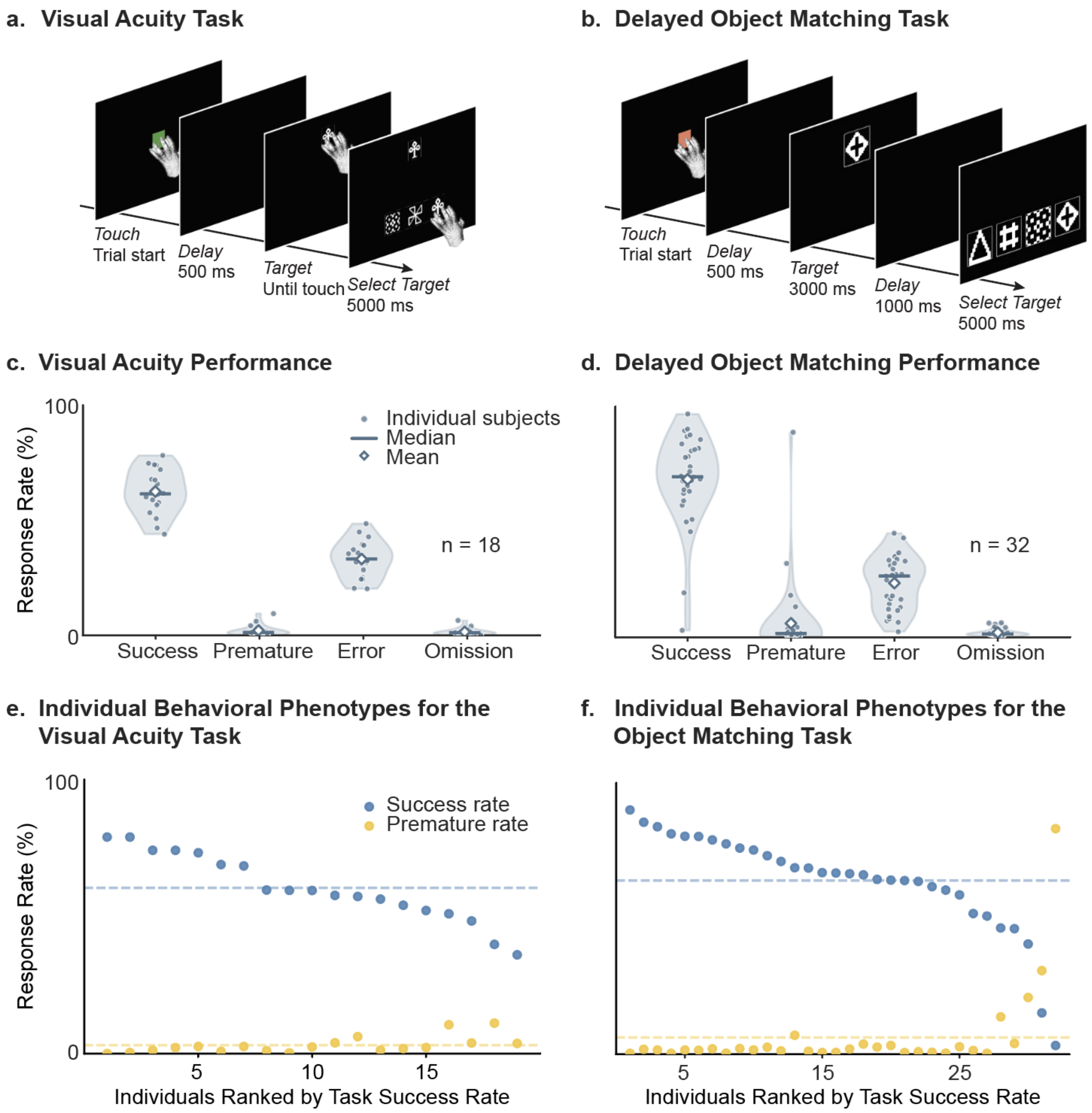


***Figure S3. Control tasks.*** *Schematic overviews of* ***(a)*** *the Visual Acuity task, assessing basic visual perception through shape discrimination with reduced-size stimuli, and* ***(b)*** *the Delayed Object Matching task, assessing visual working memory and distractor suppression through delayed stimulus matching. Distribution of success, premature, error, and omission rates across individuals in* ***(c)*** *the Visual Acuity task and* ***(d)*** *the Delayed Object Matching task. Individual behavioral phenotypes for* ***(e)*** *the Visual Acuity task and* ***(f)*** *the Delayed Object Matching task, showing mean success and premature response rates ranked by success rate; dashed lines indicate group means. As in the sustained-attention task, the lowest-ranked individuals in* ***(f)*** *show markedly elevated premature responding, indicating that impulsive responding was more evident in tasks with a memory or attentional-control demand rather than to perceptual discrimination.*

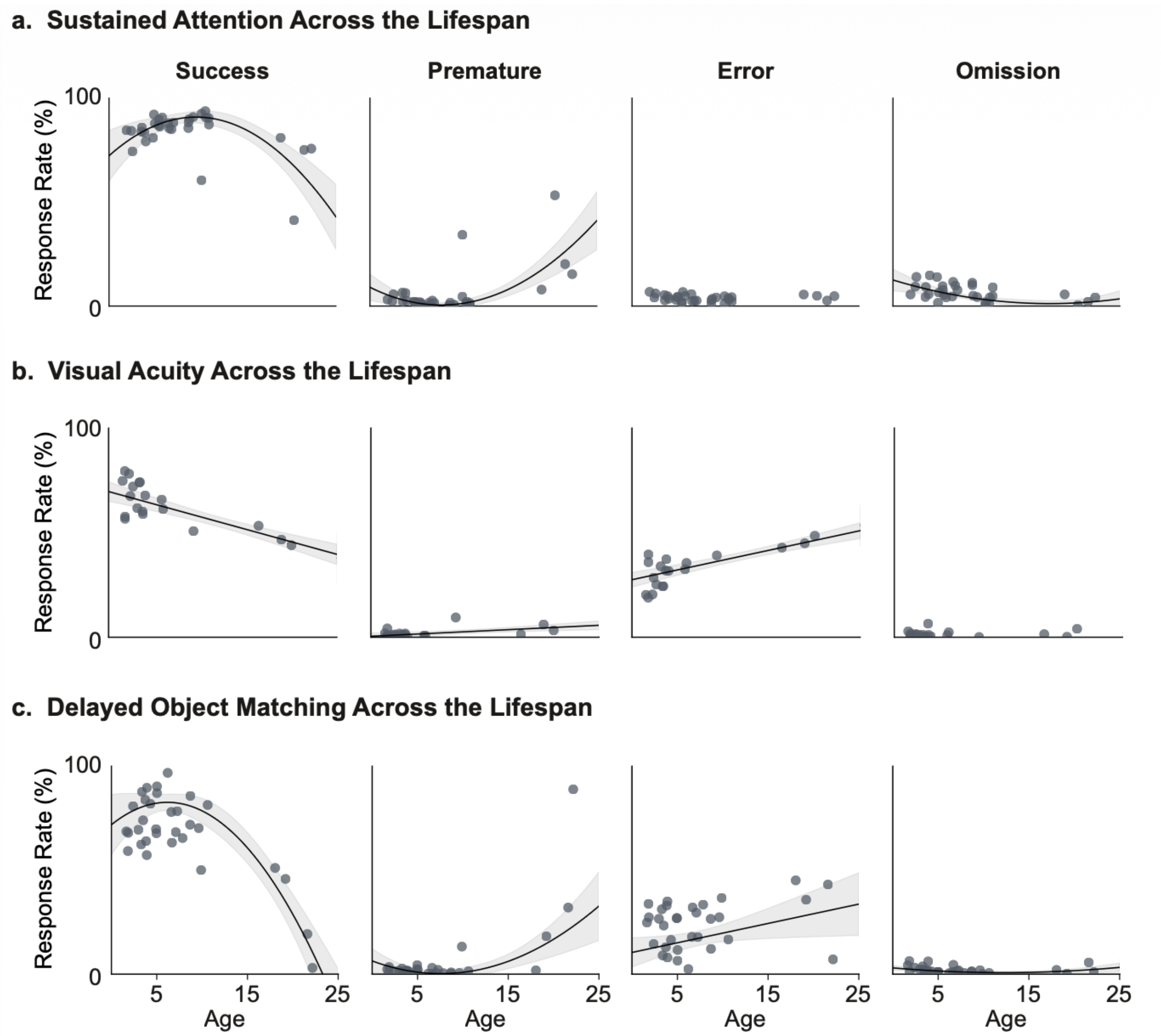


***Figure S4. Lifespan trajectories of task performance across cognitive and sensory domains.*** *Each panel shows individual animal performance estimates (dots) plotted against age at testing (years). Regression lines and 95% confidence intervals are shown only for the selected model (see Methods and* ***Table S1****).* ***(a)*** *5-Choice Serial Reaction Time Task.* ***(b)*** *Visual Acuity Task.* ***(c)*** *Delayed Object Matching Task.*

| | Success | Premature | Error | Omission |
|---|---|---|---|---|
| Model | Adjusted $R^2$ / Model p-value | | | |
| **a. Attention lifespan** - related to figure 3a | | | | |
| Linear | 0.184 / **0.008** | 0.159 / **0.014** | −0.028 / 0.700 | 0.261 / **0.002** |
| Quadratic | 0.515 / **<0.001** | 0.509 / **<0.001** | −0.024 / 0.298 | 0.362 / **0.023** |
| Linear vs. Quadratic | **<0.001** | **0.0162** | — | **0.023** |
| Selected model | Quadratic | Quadratic | — | Quadratic |
| **b. Vision lifespan** - related to figure S4b | | | | |
| Linear | 0.765 / **<0.001** | 0.390 / **0.003** | 0.771 / **<0.001** | 0.085 / 0.121 |
| Quadratic | 0.751 / 0.777 | 0.435 / 0.145 | 0.763 / 0.494 | 0.028 / 0.951 |
| Linear vs. Quadratic | — | — | — | — |
| Selected model | Linear | Linear | Linear | — |
| **c. Control lifespan** - related to figure S4c | | | | |
| Linear | 0.632 / **<0.001** | 0.074 / 0.073 | 0.120 / **0.029** | −0.006 / 0.374 |
| Quadratic | 0.785 / **<0.001** | 0.301 / **0.003** | 0.090 / 0.914 | 0.188 / **0.008** |
| Linear vs. Quadratic | **<0.001** | — | — | — |
| Selected model | Quadratic | Quadratic | Linear | Quadratic |

***Table S1. Linear and quadratic model comparisons for lifespan performance trajectories***. *For each response type (success, premature, error, omission) and task, linear and quadratic regression models were fitted to each animal's mean response rate as a function of age. Adjusted $R^2$ values quantify model goodness-of-fit. P-values correspond to the significance of the age term in the linear model ($\beta_{age}$) or the quadratic age term in the quadratic model ($\beta_{age^2}$). Model selection followed the procedure described in the Methods: when only one model showed a significant age effect (α = 0.05), that model was retained; when both models were significant, they were compared using a nested F-test, and the quadratic model was retained only if it provided a significantly better fit. Dashes in the Linear vs. Quadratic row indicate no F-test was performed; dashes in the Selected model row indicate neither model showed a significant age effect, so no regression line is shown in* ***Figures 3*** *and* ***S4***.

| | **Success** | **Premature** | **Error** | **Omission** |
|---|---|---|---|---|
| | *β (p)* | *β (p)* | *β (p)* | *β (p)* |
| *Age* | −0.015 (< .001) | 0.180 (< .001) | −0.041 (< .001) | −0.049 (< .001) |
| $Age^2$ | −0.008 (< .001) | 0.004 (< .001) | 0.005 (< .001) | −0.003 (< .001) |

***Table S2. GEE Model estimates of linear and quadratic age effects on task performance.*** *Values represent regression coefficients (β) estimated using generalized estimating equations, with age mean-centered before squaring so that the linear term reflects the slope at the average age of the sample. Significance reflects Benjamini–Hochberg FDR correction applied across all 8 tests within the task. N = 32 individuals, 641,607 trials.*

| | **Success** | **Premature** | **Error** | **Omission** |
|---|---|---|---|---|
| | *β (p)* | *β (p)* | *β (p)* | *β (p)* |
| *Social Status* | 0.153 (< .001) | −0.322 (< .001) | −0.297 (< .001) | 0.082 (< .001) |
| *Age* | −0.094 (< .001) | 0.228 (< .001) | 0.017 (< .001) | −0.084 (< .001) |
| *Social Status × Age* | −0.029 (< .001) | 0.056 (< .001) | 0.028 (< .001) | −0.057 (< .001) |

***Table S3. GEE Model estimates of social status, age, and their interaction on task performance.*** *Values represent regression coefficients (β) estimated using generalized estimating equations, with age mean-centered so that the social status coefficient reflects its effect at the sample mean age. Social status was quantified using standardized Elo ratings (M = 0, SD = 1). Significance reflects Benjamini–Hochberg FDR correction applied across all 12 tests within the model. N = 32 individuals, 641,607 trials.*

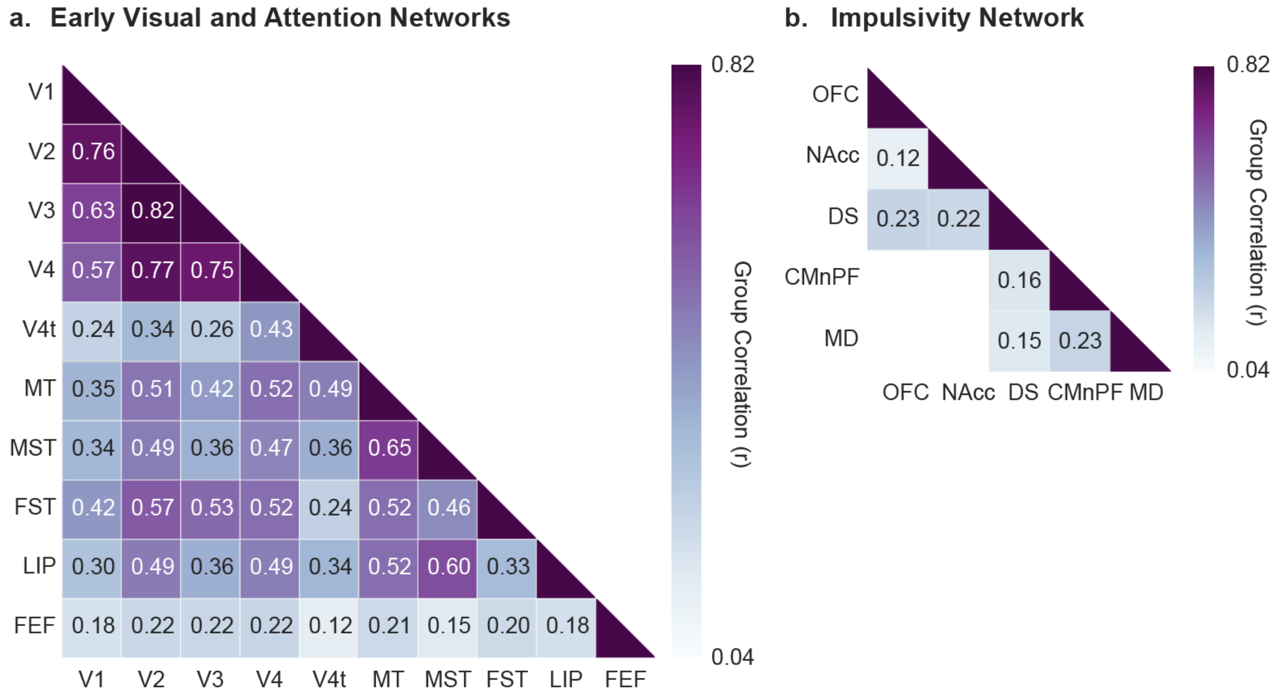


***Figure S5: Group-Level Resting-State Functional Connectivity in the Early Visual, Attentional and Cortico-Striato-Thalamic Networks.*** *Heatmaps showing mean pairwise functional connectivity (Fisher z-transformed Pearson correlations, back-transformed to r) across 18 M. tonkeana. Values shown only for connections surviving Benjamini–Hochberg FDR correction (q < 0.05) across all pairwise tests.* ***(a)*** *V1–V4: primary–quaternary visual cortex; V4t: transitional visual area 4; MT: middle temporal area; MST: medial superior temporal area; FST: fundus of the superior temporal area; LIP: lateral intraparietal area; FEF: frontal eye field.* ***(b)*** *OFC: orbitofrontal cortex; NAcc: nucleus accumbens; DS: dorsal striatum; CMnPF: centromedian-parafascicular thalamic complex; MD: mediodorsal thalamus.*

| *Attentional network* | *Linear*<br>*Adj. $R^2$ / p* | *Quadratic*<br>*Adj. $R^2$ / p* | *Lin vs. Quad*<br>*p* | *Model* |
|---|---|---|---|---|
| ***LIP–FEF*** | **-0.062 / 0.948** | **0.387 / 0.010*** | **0.003**** | **Ouadratic** |
| *LIP–V1* | 0.246 / 0.021* | 0.220 / 0.061 | — | Linear |
| *LIP–V2* | 0.111 / 0.096 | 0.226 / 0.057 | — | — |
| *LIP–V3* | -0.014 / 0.395 | 0.113 / 0.159 | — | — |
| *LIP–V4* | 0.098 / 0.110 | 0.206 / 0.069 | — | — |
| *LIP–V4t* | 0.299 / 0.011* | 0.254 / 0.044* | 0.884 | Linear |
| *LIP–MT* | 0.277 / 0.015* | 0.265 / 0.039* | 0.403 | Linear |
| *LIP–MST* | 0.488 / <0.001*** | 0.454 / 0.004** | 0.936 | Linear |
| *LIP–FST* | -0.057 / 0.780 | 0.107 / 0.168 | — | — |
| *FEF–V1* | 0.186 / 0.042* | 0.282 / 0.033* | 0.097 | Linear |
| *FEF–V2* | -0.029 / 0.480 | 0.069 / 0.229 | — | — |
| *FEF–V3* | -0.057 / 0.782 | 0.111 / 0.162 | — | — |
| *FEF–V4* | -0.057 / 0.780 | 0.238 / 0.051 | — | — |
| *FEF–V4t* | -0.053 / 0.713 | -0.023 / 0.464 | — | — |
| *FEF–MT* | -0.062 / 0.927 | 0.163 / 0.103 | — | — |
| *FEF–MST* | -0.009 / 0.370 | 0.079 / 0.210 | — | — |
| *FEF–FST* | -0.057 / 0.776 | 0.227 / 0.057 | — | — |

***Table S4. Linear and Quadratic Model Comparisons for Resting-State Functional Connectivity Lifespan Trajectories in the Attention Network.*** *Model selection followed the procedure described in the Methods: linear and quadratic models were evaluated separately for a significant effect of age ($\beta_1$ for the linear model, $\beta_2$ for the quadratic model; $\alpha = 0.05$). If only one model showed a significant age effect, that model was retained. If both were significant, they were compared using a nested F-test, and the quadratic model was retained only if it provided a significantly better fit than the linear model; otherwise the linear model was retained as more parsimonious. If neither was significant, no model was retained (—). p: p-value of $\beta_1$ (linear model) or $\beta_2$ (quadratic model). *$p < 0.05$, **$p < 0.01$, ***$p < 0.001$.*

| ***Vision network*** | ***Linear*** *Adj. $R^2$ / p* | ***Quadratic*** *Adj. $R^2$ / p* | ***Lin vs. Quad*** *p* | ***Model*** |
|---|---|---|---|---|
| *V1–V2* | 0.663 / <0.001*** | 0.640 / <0.001*** | 0.983 | Linear |
| *V1–V3* | 0.447 / 0.001** | 0.431 / 0.006** | 0.471 | Linear |
| *V1–V4* | 0.231 / 0.025* | 0.241 / 0.049* | 0.286 | Linear |
| *V1–V4t* | 0.205 / 0.034* | 0.163 / 0.103 | — | Linear |
| *V1–MT* | 0.162 / 0.055 | 0.110 / 0.163 | — | — |
| *V1–MST* | 0.100 / 0.109 | 0.091 / 0.192 | — | — |
| *V1–FST* | 0.414 / 0.002** | 0.376 / 0.011* | 0.907 | Linear |
| *V2–V3* | 0.351 / 0.006** | 0.315 / 0.023* | 0.681 | Linear |
| *V2–V4* | 0.402 / 0.003** | 0.393 / 0.009** | 0.398 | Linear |
| *V2–V4t* | -0.050 / 0.670 | -0.109 / 0.852 | — | — |
| *V2–MT* | -0.021 / 0.433 | 0.006 / 0.375 | — | — |
| *V2–MST* | -0.008 / 0.366 | 0.020 / 0.337 | — | — |
| *V2–FST* | 0.184 / 0.043* | 0.276 / 0.035* | 0.101 | Linear |
| *V3–V4* | 0.403 / 0.003** | 0.398 / 0.009** | 0.372 | Linear |
| *V3–V4t* | 0.079 / 0.136 | 0.175 / 0.093 | — | — |
| *V3–MT* | 0.063 / 0.163 | 0.354 / 0.015* | — | Quadratic |
| *V3–MST* | -0.059 / 0.833 | 0.219 / 0.061 | — | — |
| *V3–FST* | 0.256 / 0.019* | 0.280 / 0.033* | 0.232 | Linear |
| *V4–V4t* | -0.043 / 0.588 | -0.094 / 0.768 | — | — |
| *V4–MT* | -0.010 / 0.375 | 0.059 / 0.248 | — | — |
| *V4–MST* | -0.060 / 0.846 | 0.055 / 0.256 | — | — |
| *V4–FST* | 0.123 / 0.085 | 0.333 / 0.019* | — | Quadratic |
| *V4t–MT* | -0.031 / 0.493 | -0.099 / 0.796 | — | — |
| *V4t–MST* | 0.294 / 0.012* | 0.275 / 0.035* | 0.453 | Linear |
| *V4t–FST* | -0.043 / 0.595 | -0.068 / 0.639 | — | — |
| *MT–MST* | 0.308 / 0.010** | 0.265 / 0.039* | 0.810 | Linear |
| *MT–FST* | 0.008 / 0.302 | 0.202 / 0.072 | — | — |
| *MST–FST* | -0.062 / 0.997 | 0.222 / 0.059 | — | — |

***Table S5. Linear and Quadratic Model Comparisons for Resting-State Functional Connectivity Lifespan Trajectories in the Visual Network.*** *Model selection followed the procedure described in the Methods: linear and quadratic models were evaluated separately for a significant effect of age ($\beta_1$ for the linear model, $\beta_2$ for the quadratic model; $\alpha = 0.05$). If only one model showed a significant age effect, that model was retained. If both were significant, they were compared using a nested F-test, and the quadratic model was retained only if it provided a significantly better fit than the linear model; otherwise the linear model was retained as more parsimonious. If neither was significant, no model was retained (—). p: p-value of $\beta_1$ (linear model) or $\beta_2$ (quadratic model). *$p < 0.05$, **$p < 0.01$, ***$p < 0.001$.*

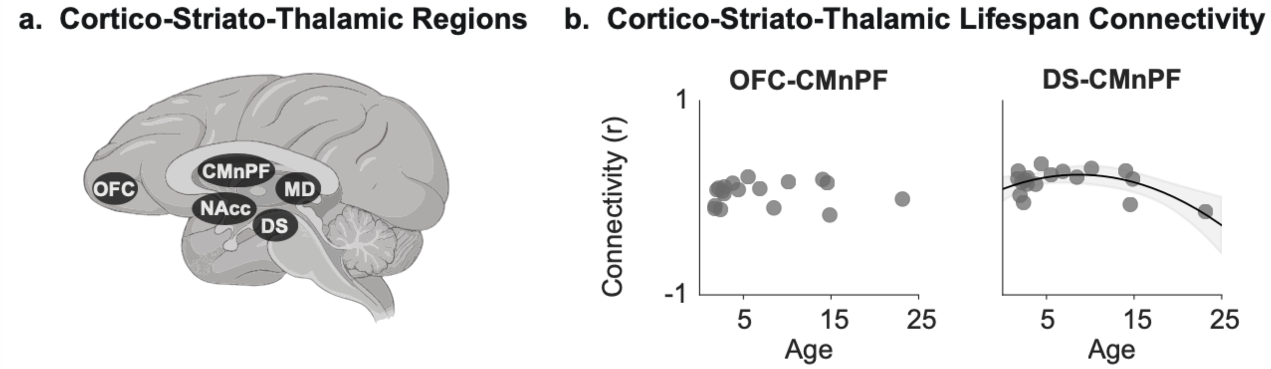


***Figure S6. Regions of Interest and Lifespan Trajectories Along Cortico-Striato-Thalamic Connections. (a)*** *Regions of interest.* ***(b)*** *Lifespan functional connectivity of the Cortico-Striato-Thalamic network: Fisher z-transformed connectivity plotted as a function of age at scan, shown for OFC–CMnPF and DS–CMnPF connectivity. DS–CMnPF showed a significant quadratic trajectory (regression line and 95% CI shown); all other connections, including OFC–CMnPF, remained stable across the lifespan. CMnPF: centromedian-parafascicular thalamic complex; DS: dorsal striatum; OFC: orbitofrontal cortex.*

| ***Cortico-striato-thalamic connections*** | ***Linear*** *Adj. $R^2$ / p* | ***Quadratic*** *Adj. $R^2$ / p* | ***Lin vs. Quad*** *p* | ***Model*** |
|---|---|---|---|---|
| *MD–DS* | -0.062 / 0.950 | -0.133 / 0.997 | — | — |
| *MD–NAcc* | -0.044 / 0.597 | -0.032 / 0.495 | — | — |
| *MD–OFC* | -0.054 / 0.726 | -0.124 / 0.941 | — | — |
| *MD–CMnPF* | -0.052 / 0.700 | -0.107 / 0.841 | — | — |
| *DS–NAcc* | -0.059 / 0.813 | -0.124 / 0.941 | — | — |
| *DS–OFC* | -0.049 / 0.652 | -0.079 / 0.691 | — | — |
| *DS–CMnPF* | 0.064 / 0.160 | 0.348 / 0.016* | — | Quadratic |
| *NAcc–OFC* | -0.062 / 0.962 | -0.130 / 0.977 | — | — |
| *NAcc–CMnPF* | -0.060 / 0.838 | 0.029 / 0.314 | — | — |
| *OFC–CMnPF* | -0.062 / 0.989 | -0.043 / 0.536 | — | — |

***Table S6. Linear and Quadratic Model Comparisons for Resting-State Functional Connectivity Lifespan Trajectories Across Cortico-Striato-Thalamic Connections.*** *Model selection followed the procedure described in the Methods: linear and quadratic models were evaluated separately for a significant effect of age ($\beta_1$ for the linear model, $\beta_2$ for the quadratic model; $\alpha = 0.05$). If only one model showed a significant age effect, that model was retained. If both were significant, they were compared using a nested F-test, and the quadratic model was retained only if it provided a significantly better fit than the linear model; otherwise the linear model was retained as more parsimonious. If neither was significant, no model was retained (—). p: p-value of $\beta_1$ (linear model) or $\beta_2$ (quadratic model). *$p < 0.05$, **$p < 0.01$, ***$p < 0.001$.*

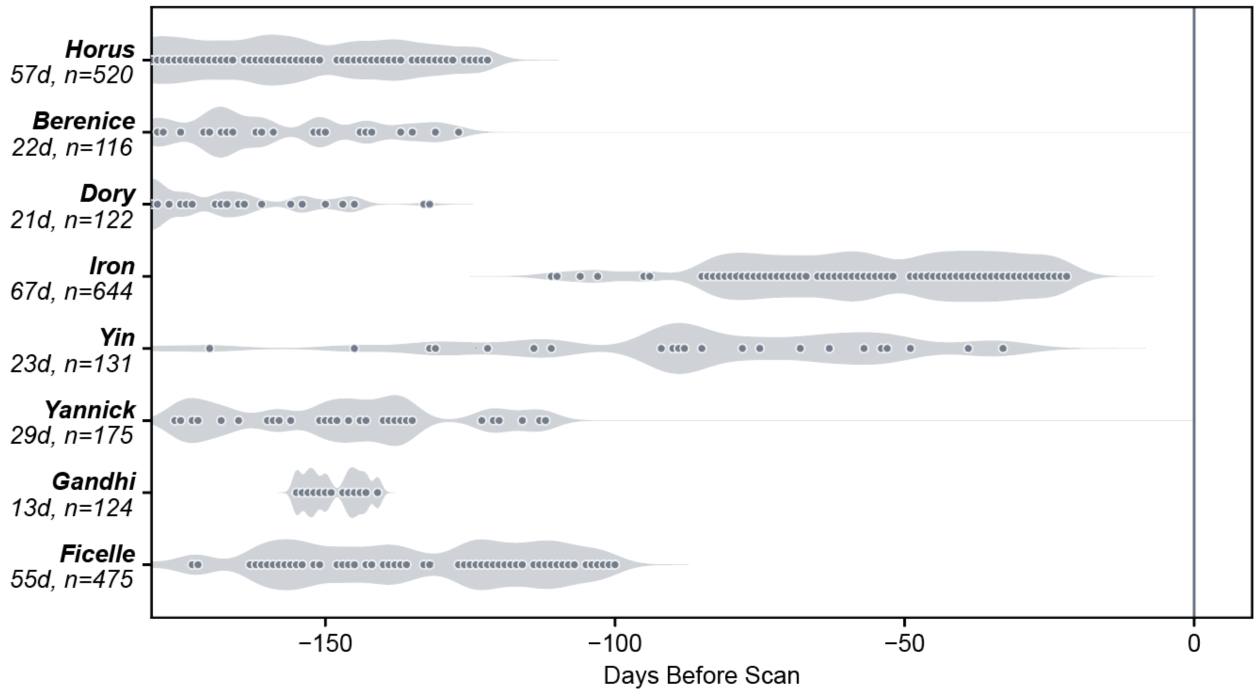


***Figure S7: Behavioral Testing Activity During the 6 Months Preceding Neuroimaging.*** *Timeline of behavioral testing activity prior to neuroimaging. Each dot marks a day when a subject performed the task during the 6-month pre-scan window. The shaded violin shape shows the density of testing activity over time, with wider regions indicating periods of more frequent task engagement (weighted by daily attempt counts). The vertical line marks the scan date (day 0); negative x-axis values indicate days before scanning. Subject labels show total testing days (d) and total task trials (n) within the 6-month window.*

| | *Success* r (p) | *Premature* r (p) | *Omission* r (p) | *Error* r (p) |
|---|---|---|---|---|
| ***Visuo-attentional network*** | | | | |
| ***LIP–FEF*** | **0.833 (0.010)** | **—** | **-0.842 (0.009)** | **—** |
| *LIP–V2* | — | — | -0.748 (0.033) | — |
| *LIP–V4* | — | — | -0.746 (0.034) | — |
| *LIP–V4t* | — | — | -0.817 (0.013) | — |
| *LIP–MT* | — | 0.714 (0.047) | — | — |
| *FEF–V1* | 0.880 (0.004) | — | -0.756 (0.030) | — |
| *FEF–V2* | 0.819 (0.013) | — | — | — |
| *FEF–V3* | 0.831 (0.011) | — | — | — |
| *FEF–V4* | 0.925 (0.001) | — | — | — |
| ***Visuo-visual networks*** | | | | |
| *V1–V3* | — | — | — | -0.811 (0.015) |
| *V2–V3* | — | — | — | -0.737 (0.037) |
| *V2–V4t* | — | — | — | -0.908 (0.002) |
| *V3–V4t* | — | — | — | -0.842 (0.009) |
| *V4–V4t* | — | — | — | -0.713 (0.047) |
| *V4–MST* | — | — | 0.741 (0.035) | — |
| *V4t–FST* | — | — | — | -0.901 (0.002) |
| *V4t–MT* | — | — | — | -0.877 (0.004) |
| *V4t–MST* | — | — | — | -0.723 (0.043) |

***Table S7. Age-Adjusted Brain–Behavior Correlations Across Attention and Visuo-Visual Networks.*** *Age-adjusted partial correlations controlling for age at scan (df = 5, n = 8); p-values derived from t-test on the partial correlation coefficient. Only connections with at least one significant association (uncorrected $p < 0.05$) are shown. No associations survived FDR correction (Benjamini–Hochberg, $q < 0.05$) across 180 tests (45 connections × 4 outcomes); results are therefore interpreted as exploratory. FEF: frontal eye field; LIP: lateral intraparietal area; MT: middle temporal area; MST: medial superior temporal area; FST: fundus of the superior temporal area; V1–V4t: visual areas 1–4 and transitional area 4.*

| | *Success* | *Premature* | *Omission* | *Error* |
|---|---|---|---|---|
| | *r (p)* | *r (p)* | *r (p)* | *r (p)* |
| ***Cortico-striato-thalamic connections*** | | | | |
| *OFC–CMnPF* | 0.853 (0.007) | — | -0.940 (0.001)* | — |
| *OFC–NAcc* | — | -0.763 (0.028) | 0.737 (0.037) | — |
| *OFC–MD* | — | 0.821 (0.013) | — | — |
| *DS–CMnPF* | — | -0.798 (0.018) | — | — |
| *DS–NAcc* | — | — | 0.743 (0.035) | — |
| *MD–CMnPF* | -0.717 (0.045) | — | — | — |

***Table S8. Age-Adjusted Brain–Behavior Correlations Across Cortico-Striato-Thalamic Connections.*** *Age-adjusted partial correlations controlling for age at scan (df = 5, n = 8); p-values derived from a t-test on the partial correlation coefficient. Only connections with at least one significant association (uncorrected p < 0.05) are shown. FDR correction (Benjamini–Hochberg, q < 0.05) was applied within this network (40 tests: 10 connections × 4 outcomes). One association survived correction: OFC–CMnPF connectivity was negatively associated with omission rate (r = -0.940, uncorrected p = 0.0005, FDR-adjusted p = 0.020). All other associations are uncorrected and interpreted as exploratory. OFC: orbitofrontal cortex; NAcc: nucleus accumbens; DS: dorsal striatum; CMnPF: centromedian-parafascicular thalamic complex; MD: mediodorsal thalamus.*
